# Evolution of parasitism-related traits in nematodes

**DOI:** 10.1101/2025.09.26.678730

**Authors:** Chieh-Hsiang Tan, Hillel T. Schwartz, Nathan Y. Rodak, Paul W Sternberg

## Abstract

The abundant resources provided by the host provide an evolutionary rationale for parasitism and drive the metabolic and developmental divergence of parasitic and free-living animals. Two evolutionarily distant nematode genera, *Steinernema* and *Heterorhabditis*, independently evolved an entomopathogenic lifestyle, in which they invade insects and kill them with the assistance of specifically associated symbiotic pathogenic bacteria. It had been generally assumed that the worms, being bacterivores, feed on their symbiotic bacteria, which rapidly reproduce as they consume the insect host. The evolutionary adaptations of entomopathogenic nematodes to a parasitic lifestyle developmentally, and the symbiotic relationships of entomopathogenicity, remain largely unknown. We developed an axenic culture medium that supports robust and sustained growth of *Steinernema hermaphroditum*, with finite control of nutrients available to the nematodes. We found that, uniquely among the nematodes tested, *S. hermaphroditum* hatchlings cannot endure in a nutrient-poor environment; this ability is impaired but still present in *Heterorhabditis bacteriophora.* Similarly, the ability to forage for food is completely lost in *H. bacteriophora* hatchlings and severely compromised in *S. hermaphroditum*. We reasoned that these traits were lost because they are unnecessary to obligate parasites that always hatch in a resource-rich host. We further found that *Steinernema* and, to a limited extent, *Heterorhabditis* nematodes can successfully invade, develop, and reproduce inside a living insect host independent of their symbiotic bacteria, apparently feeding on the hemolymph, and emerge carrying bacteria found within, explaining the evolutionary origins of entomopathogenic nematodes.

**Highlights:**

- A simple but robust axenic culturing method for the emerging model nematode *Steinernema hermaphroditum* and other nematode parasites of invertebrates.
- Nematode adaptation to parasitism is associated with changes in modes of feeding.
- Convergent evolution led to the loss of hatchling survival traits in entomopathogenic nematodes.
- Entomopathogenic nematodes evolved from parasitoid ancestors and likely acquired their bacterial symbionts from their hosts.

**Graphical abstract:** 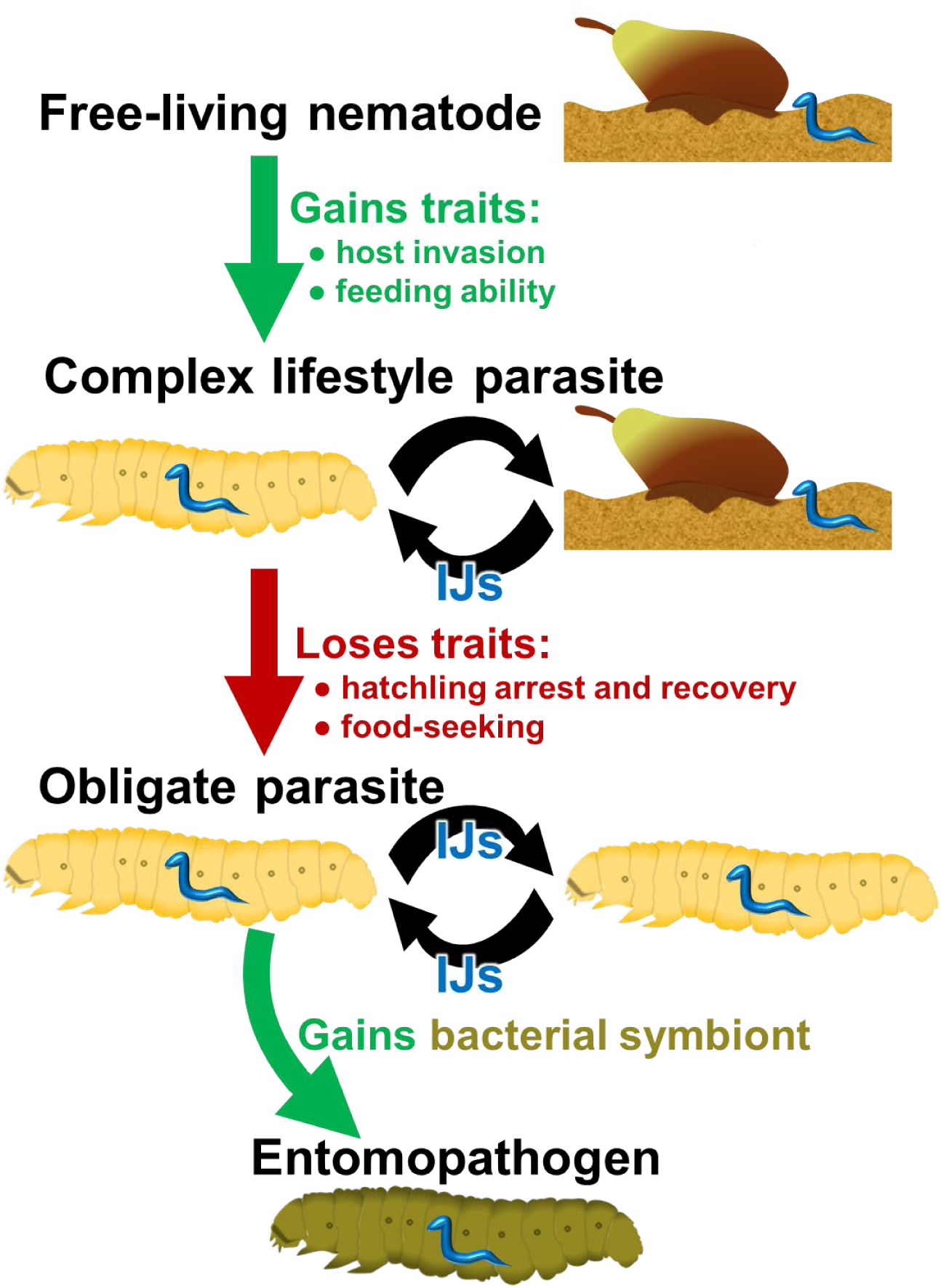

## Introduction

Each living organism is a concentration of the resources necessary for life and therefore constitutes an ecological opportunity that is bound to be exploited. Parasites, in a multitude of iterations and variations, have frequently evolved to fill this niche. In adapting to the resource-rich environment of a host, an obligate parasite should, in theory, lose unnecessary traits that helped it survive in other, resource-poor environments, now left behind, and develop traits that enable it to utilize the host’s rich resources. Many examples can be found of parasites that have lost the ability to produce some molecules that they can more easily obtain from their host.^1–3^ Examples of parasites adapting to the environment of their hosts include mechanisms of immune evasion,^4–6^ adaptations in their feeding mechanisms,^7,8^ or alterations in how they metabolize the nutrients they acquire.^9^ The loss of traits unnecessary to a host organism and the development of traits specifically useful to a host should explain most of the developmental and physiological divergences between parasites and their free-living relatives.

As perhaps the most common metazoan, nematodes occupy nearly every ecological niche in our ecosphere.^10,11^ Small, mostly transparent, often featuring rapid growth and short generation times, and in many cases readily maintained in the laboratory, these animals offer fertile ground for the observation of their development and description of their evolutionary adaptations, and for experimental interventions to explore the mechanisms underlying these processes. Parasitism has arisen repeatedly and independently throughout nematode evolution, with parasitic species nested within free-living species phylogenetically.^12^ Some of these nematodes have evolved remarkably similar lifestyles, as seen in *Steinernema* and *Heterorhabditis*, which are situated on branches of the nematode phylogeny that likely separated more than 200 million years ago.^13,14^ The *Steinernema* and *Heterorhabditis* genera each consist of dozens of species, all of which are entomopathogenic nematodes (EPNs),^15^ which are defined by: (1) their ability to infect insects as infective juvenile (IJ) larvae, an environmentally resistant developmentally arrested alternate larval stage equivalent to the dauer larvae of *Caenorhabditis elegans* and other free-living nematodes; (2) their association with dedicated symbiotic bacteria, which they carry within the IJ protected from the environment, and (3) the rapidity and efficiency with which they and their bacteria kill and consume their insect prey. The bacteria use the carcass as a substrate for reproduction, while inhibiting the growth of other microorganisms and deterring predators and scavengers other than their nematode partners. The resulting bacterial monoculture serves as a food source well suited to the needs of their nematode partners.^16–18^

The origins of the entomopathogenic lifestyle are unclear, and it is presumed to have arisen from one of the other existing modes of interaction between nematodes and insects: phoresis, necromeny, entoecy, or parasitism.^19,20^ Nematodes that have a phoretic relationship with insects are attracted to the insects and use them as a means of conveyance, departing the insect when it arrives at a suitable environment for their growth. Necromenic nematodes wait on the insect for its natural death, then resume development and feed on its carcass. Entoecic nematodes feed on the bacteria and fungi growing in or on their host, without feeding on their host. Parasitic nematodes invade the host’s body and directly feed on the living animal. Remnant capacity for any of these lifestyles in entomopathogenic nematodes could provide an indication of their evolutionary origins.

One of the most striking differences in nematode lifestyle is between free-living animals and obligate parasites.^21^ Free-living animals hatch into a complex environment that they must explore to find the best available food source and escape from predators; parasites hatching in their host have their needs provided for; if, for some reason, they do not hatch within a viable host, they have no possibility of reaching another. This means that in free-living nematodes and in parasites with a complex life cycle involving a free-living stage, but not in obligate parasites, there should be evolutionary pressure to maintain the ability of a newly hatched nematode to survive temporarily adverse conditions and to maintain its tendency to explore its environment in search of a suitable food source.

Here, we report our observations regarding the parasitic origins and their evolutionary consequences for *Steinernema hermaphroditum* and its fellow entomopathogen, *Heterorhabditis bacteriophora*. We developed a method for the axenic culture of *S. hermaphroditum* and found that *S. hermaphroditum* shares with other parasites of invertebrates a feature absent in numerous representatives of free-living nematodes: the ability to grow rapidly and reproduce when fed only on minimally supplemented bacterial growth media. We find that, as they evolved a dedicated entomopathogenic lifestyle, both *Steinernema* and *Heterorhabditis* independently and differently lost the ability of newly hatched larval nematodes to survive in the absence of immediately available food and to locate a suitable food source. We demonstrate that when the EPNs *Heterorhabditis* and *Steinernema* are deprived of their bacterial symbiont, and so are unable to rapidly kill insects and lack their preferred food source, they can feed directly upon the insect to develop to reproductive adulthood; *Steinernema* in particular are able to grow essentially as quickly as they would grow in insects killed by their bacterial symbiont, and to the same size. The infective juveniles (IJs) produced by these asymbiotic infections emerge carrying bacteria picked up from their insect host, providing a clear mechanism by which their existing symbiotic partnership could have arisen.

## Results

### Simple, scalable axenic culture of *Steinernema hermaphroditum* for developmental studies and the mass production of infective juveniles

To investigate the developmental and metabolic adaptations of EPNs and the origin of the complex triangular symbiotic relationship of entomopathogenicity, we developed an axenic culture medium that allows for robust and sustained growth of *Steinernema hermaphroditum,* a worm that we have been developing as a model for such studies.^13,22–26^

The medium has three major components: (1) as its primary carbon source, tryptone and yeast extract, in the same relative proportion as in the bacterial growth medium LB;^27^ (2) a trace metal solution used in the liquid culture of *C. elegans*;^28^ (3) hemin and cholesterol, as nematodes are auxotrophs for both heme and sterols (Figure 1A). ^28,29^ All three major components were necessary for sustained culture, and the concentration of the carbon source ingredients was titrated based on the growth and reproduction of *S. hermaphroditum*, which we found to grow best between 3X and 3.5X the strength of LB, with a reduced 1X LB strength proving suitable for rapid and consistent IJ production. This axenic culture protocol was initially developed in small volumes in 6-well to 24-well plates or in 15 mL conical tubes, but was easily adapted to several hundred milliliters shaken in Erlenmeyer flasks (Figure 1B).

**Figure 1.**
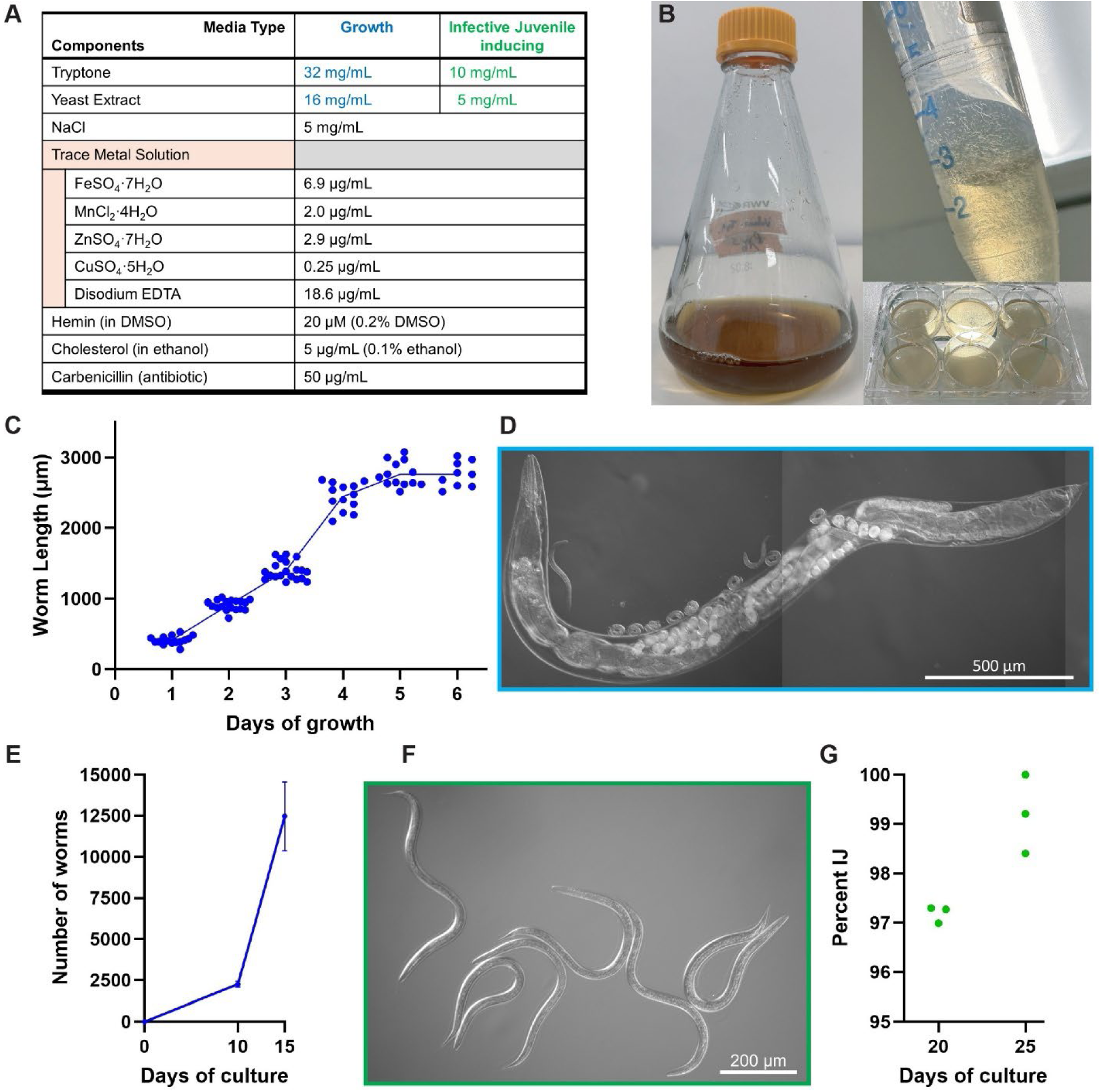
A robust, simple, and scalable axenic culture of *Steinernema hermaphroditum*. (A) The compositions of the axenic culture media used in this study. The top two rows are divided into two columns: the left column (Growth) is a media composition optimized for growth and reproduction, while the right column (Infective Juvenile Inducing) is a composition optimized for IJ production. (B) The media is clear and easily scalable, from tissue culture plates and 15-ml tubes to flasks holding hundreds of milliliters (as shown). (C-D) Growth in axenic media sustained stable development of *S. hermaphroditum*, similar to that of bacterial-based liquid culture (see Figures S1A and S1B). (C) The lengths of worms were measured each day after embryos were placed in the media (n= 9-20). (D) An adult worm from the axenic culture, 5 days after eggs were placed in media. Due to the lack of pigmentation from bacteria, the worm is highly transparent. Scale bar: 500 µm. (E) Worms reproduce exponentially in this medium. Numbers were calculated based on three samples removed from the same 200mL culture at each time point. Eggs were seeded at day 0 at a density estimated to be 4.4 ± 0.5 per mL (viable) (n=3). (F-G) IJs can be effectively produced with the “Infective Juvenile inducing” formula given in part (A). (F) Infective juveniles from 20-day culture. The worms were completely enclosed by an unshed second larval stage (J2) cuticle. Scale bar: 200 µm. (G) The “Infective Juvenile inducing” formula can produce a worm population consisting almost entirely of IJs; about 97% and 99% of the population were IJs by day 20 and 25, respectively. Eggs were seeded at day 0 at a density estimated to be 185.3 ± 15.3 per mL (viable) (n=3). Data points were from three biological replicates in the same experiment. At least 250 worms were scored in each data point. In panels C and E, Values are mean ± SD.

The media is clear and transparent (Figure 1B), allowing easy observation of the cultured animal, in sharp contrast to turbid cultures using bacteria as a food source.^26^ The worms developed robustly in this medium, reaching their full size by day 5 (Figures 1C and 1D), similar to the speed and size achieved in liquid culture using a bacterial food source (Figures S1A and S1B), but slightly slower in growth and smaller in size compared to animals grown on bacterial lawns on solid agar-based media.^24^ Cultures could exhaust their available nutrients and achieve high population densities, conditions that induced the production of developmentally arrested IJs; these IJs were healthy and capable of infecting insects (see below). When grown in a sufficient volume to avoid crowding and exhaustion of nutrients, axenic cultures could be maintained for multiple generations, allowing exponential population growth (Figure 1E).

A major advantage of using this medium is the ability to precisely adjust the nutrient availability to that of the nematodes. We found that reducing the concentration of carbon sources provided to the nematodes facilitated the transition of the population into a nearly pure population of IJ larvae (>99%; Figures 1F and 1G).

### *Steinernema* hatchlings fail to developmentally arrest in the absence of food

In exploring the effects of reduced nutrient concentration, we found that hatchlings of *S. hermaphroditum* died rapidly under low nutrient conditions (Figure 2A). This is in sharp contrast to previous observations that when *C. elegans* and closely related species hatch in the absence of food, they enter a state of developmental arrest. This state allows them to conserve energy and remain viable for several weeks, resuming development when a food source is provided.^30–34^ The hatchlings of *S. hermaphroditum* normally emerge inside the resource-rich environment of the host, and if they were to hatch outside a host, they would have no way to invade a host. We hypothesized that, after *Steinernema* became entomopathogenic, it lost all evolutionary pressures to maintain an ability for hatchlings to developmentally arrest when deprived of food. The corollary was that nematodes sharing the free-living lifestyle of *C. elegans* should retain a similar ability; it was not known how widespread developmental arrest of food-deprived hatchlings is across nematodes.

**Figure 2.**
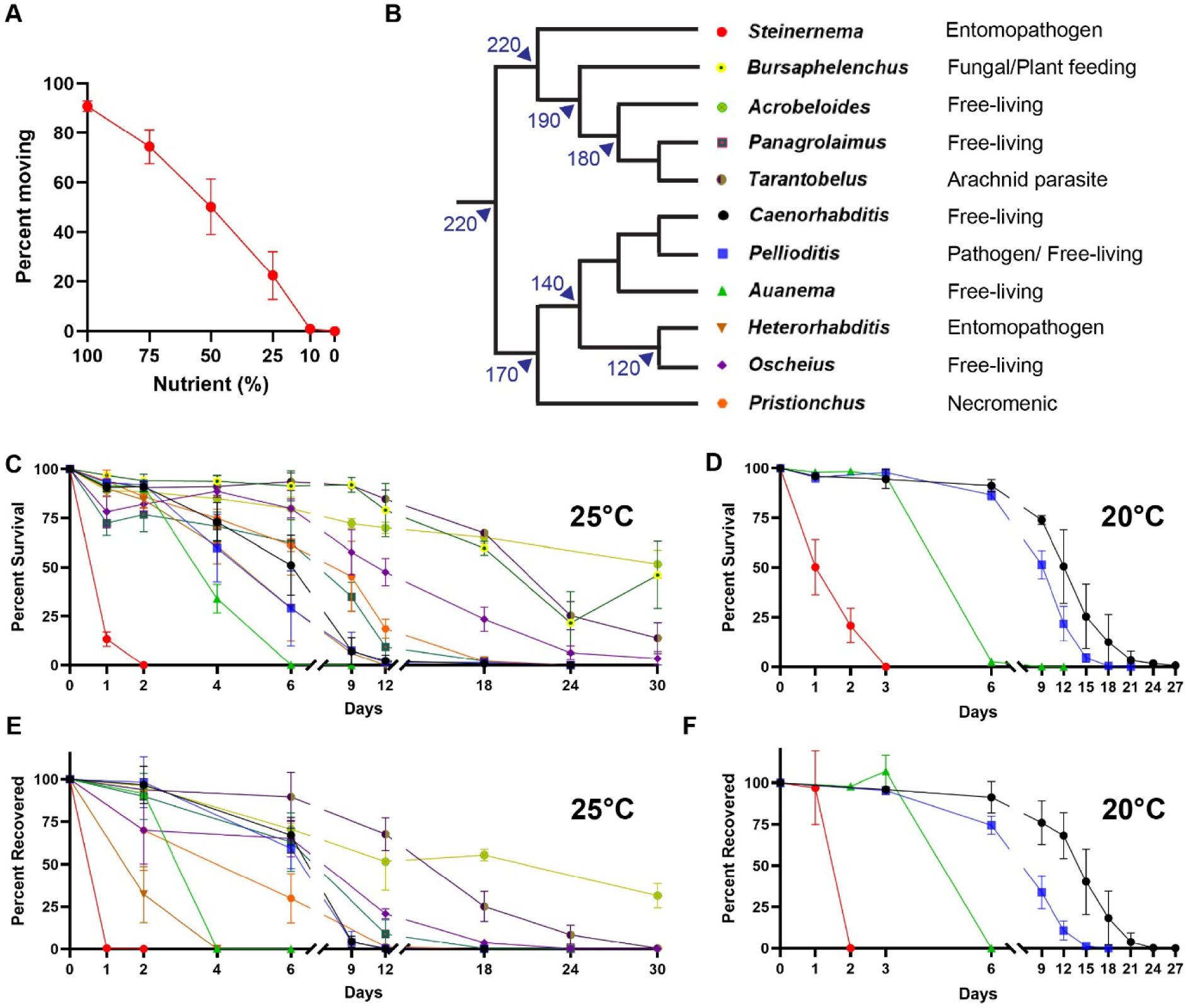
The ability to developmentally arrest is lost in *Steinernema* hatchlings. (A) *Steinernema hermaphroditum* hatchlings could not endure starvation. Eggs were hatched in axenic media containing different concentrations of carbon sources, and the proportion of animals still moving at 48 hours was scored. When nutrient levels were low, most worms ceased movement within the assay. 100% nutrients is the equivalent of the Infective Juvenile inducing medium. (n=4, 10-44 worms were assayed for each sample. See Methods for details.) (B) Phylogeny of nematode genera used in this study, with approximate evolutionary distances adapted from Schwarz *et al.* ^13^ and Qing *et al.* ^14^, normalized to the estimates given in the former, which are more conservative and approximately 1/3 lower than the latter. (C and D) Percentage of nematode hatchlings alive days after hatching into a nutrient-less environment at (C) 25°C, and (D) 20°C. *S. hermaphroditum* hatchlings are uniquely bad at enduring starvation. Every other species examined possessed at least some ability to maintain viability for at least several days. (n=3-4, see Methods for details). (E-F) Percentage of nematode hatchlings that were able to resume growth when provided with food, days after hatching into a nutrient-less environment at (E) 25°C, and (F) 20°C. *S. hermaphroditum* hatchlings endure for less than 24 hours at their normal culturing temperature (25°C) before terminally losing their viability. The hatchlings of the evolutionarily distant but ecologically similar *H. bacteriophora* endure starvation much worse than the other nematodes (See Methods for how the percentages were calculated). In panels A and C-F, Values are mean ± SD.

We therefore analyzed a broad panel of nematodes from 11 genera representing lifestyles ranging from parasitism to free-living (Figure 2B), finding that every species examined, other than *S. hermaphroditum*, possessed at least some ability to maintain viability for at least several days when hatched in the absence of food (Figures 2C and 2D). Hatchlings of some species remained visibly healthy and vigorous in nutrient-free liquid for up to several weeks, while others ceased movement within days or adopted a shrunken appearance, but some of these motionless or shrunken larvae demonstrated their viability by resuming development when placed on an appropriate food source (see Methods). We found that the appearance and activity of the starved hatchlings were lagging indicators of their eventual survival: populations lost the ability to resume growth upon introduction to food more rapidly than they lost the appearance of vitality (Figures 2E and 2F). In *S. hermaphroditum*, the ability to resume growth was abolished within 24 hours of food deprivation, measured from when their eggs were obtained and axenized. This could be extended by 24 hours if the eggs were cultured at 20°C instead of their normal 25°C, but much of this extension may be due to the additional time spent completing embryonic development at this reduced temperature. Although the hatchlings of *H. bacteriophora*, the other EPN in this study, remained active and visibly living for more than a week in the absence of food (Figure 2C), starved *H. bacteriophora* hatchlings rapidly lost their ability to resume development (Figure 2E). The hatchlings of two other parasitic nematodes, the slug pathogen *Pellioditis hermaphrodita*,^35,36^ and the arachnid parasite *Tarantobelus jeffdanielsi*,^37^ retained the ability to arrest fully. However, this is unsurprising in the case of *P. hermaphrodita*, as it is known to be a facultative parasite capable of living on feces and leaf litter,^38,39^ with closely related free-living species within its genus.^40,41^ *T. jeffdanielsi* has only recently been discovered, as one of only two described species in its genus, and it is not yet known whether they are obligate parasites or whether their life cycles might include free-living populations.

### *Heterorhabditis* hatchlings lack food-seeking behavior

Although they performed worse in assays of hatchling arrest and recovery than any other tested species besides *S. hermaphroditum*, *H. bacteriophora* hatchlings showed some limited ability in both assays (Figures 2C and 2D). This ability was unexpected, as we had found that when *H. bacteriophora* eggs hatched on a plate in the absence of food, the hatchlings invariably died, even when an ample lawn of their bacterial symbiont was only millimeters away. We examined our panel of divergent nematodes for the ability to react to hatching in the immediate absence of food by searching for a hood source. We placed their axenized eggs either on a food source (to confirm their viability) or ∼4 cm away from a food source (Figure 3A). *H. bacteriophora* hatchlings never left the site of their deposition and never found the lawn, while at least some hatchlings of every other species tested could reach the lawn within 36 hours, although *Panagrolaimus* (∼3%) and especially *Steinernema* (∼1%) hatchlings were very bad at finding their food source (Figure 3B). In the case of *Panagrolaimus*, part of this maybe due to the relatively slower hatching of the worm compare to other species, and a significantly higher proportion (∼11%) were able to reach food by 60 hours (Figure 3C); *S. hermaphroditum* hatchlings did not improve on their food-finding over time, presumably because any animals that did not reach the food source before 36 hours had elapsed would have starved to death (Figure 2E and 3C).

**Figure 3.**
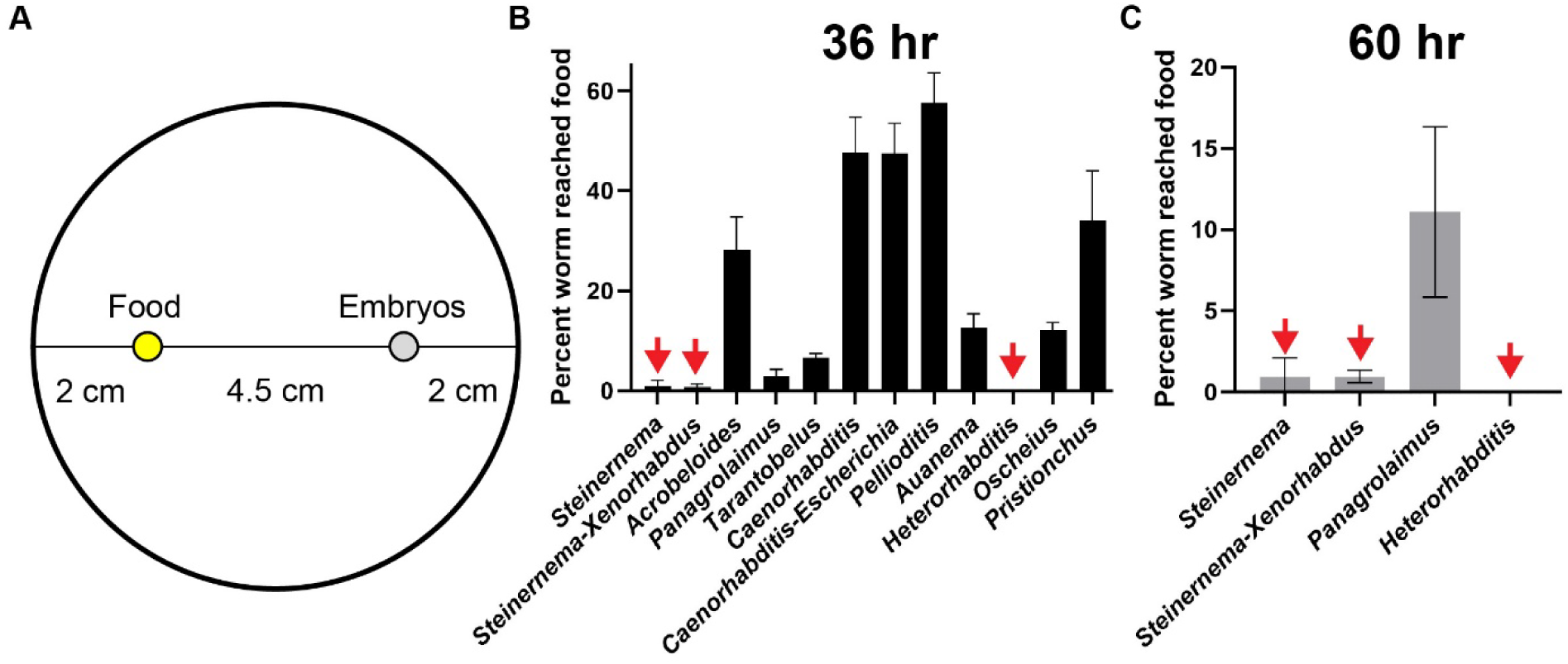
Food-seeking behavior is lost in *Heterorhabditis* hatchlings. (A) The experimental design of the food-seeking experiment. The axenized nematode embryos (grey) were seeded ∼4 cm away from their food source (yellow). *Heterorhabditis* was offered a lawn of its *Photorhabdus* bacterial symbiont; all other species were offered a lawn of *Comamonas aquatica* bacteria, with additional trials offering *Steinernema* its *Xenorhabdus* symbiont and offering *Caenorhabditis* its usual *Escherichia coli* food strain OP50. (B-C) *Heterorhabditis* and *Steinernema* hatchlings were uniquely bad at food seeking (Red arrows), with less than 1% of *Steinernema* hatchlings reaching the food 36 (B) and 60 hours (C) into the experiment. *Heterorhabditis* hatchlings completely lack food-seeking behavior; 0% of the worms ever reached their food or even left the area where the embryos were deposited. In panels B-C, Values are mean ± SD (n=4).

### Parasitic but not free-living worms thrived in the axenic culture

We tested whether our simplified axenic media could be applied to nematode culture in general. Examining our panel of divergent nematodes, we found that although many worms can grow slightly larger from hatchlings, sustained growth and reproduction were only observed in 3 of the 4 parasitic genera and not in any of the seven free-living genera (Figure 4A). In *Tarantobelus. jeffdanielsi*, growth was robust and not dissimilar to that in bacterial-based liquid culture (Figures 4B, 4C, S1C, S1D). The growth of *Pellioditis hermaphrodita* was slow and uneven, but the species could be maintained in axenic culture for multiple generations. Significant growth was also observed in *H. bacteriophora*, but their development arrested during larval development, likely due to the previously reported specific dependence of the worm on its *Photorhabdus* bacterial symbiont for growth in culture.^42–44^ Of the free-living species surveyed, only *C. elegans* produced adults in our axenic culture media; the growth of *C. elegans* was uneven, as we had seen for *P. hermaphrodita*, but unlike *P. hermaphrodita,* we were not able to continuously culture a rapidly growing population of *C. elegans*: although some F_1_s were produced, fertility was very limited. We also tested dioecious species of *Steinernema* (*carpocapsae*, *feltiae*, and *glaseri*) and *Pellioditis* (*pelhamensis*) and found significant growth but limited fertility; these species may have difficulty mating in continuously agitated liquid media.

**Figure 4.**
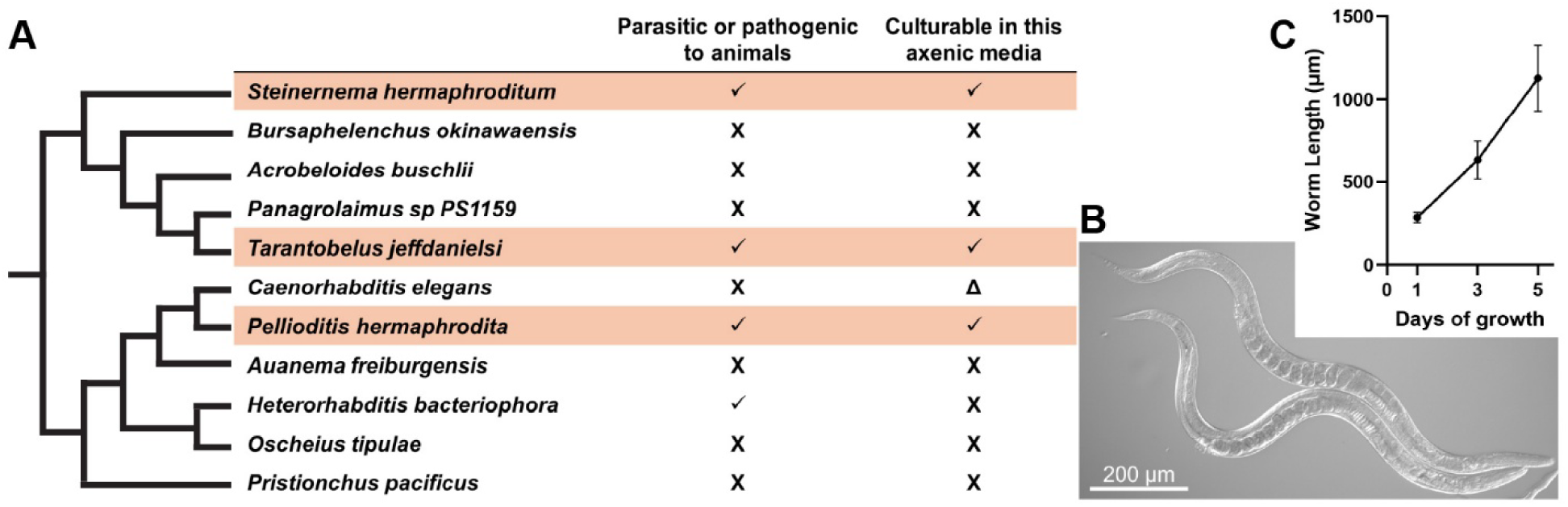
Parasitic but not free-living worms thrived in the axenic culture. (A) Among the nematodes we surveyed, we found three species that were axenically culturable in minimally supplemented bacterial growth media, all of which are parasitic (indicated by orange bars). Additionally, *Caenorhabditis elegans* sometimes reached adulthood, but could not be continuously cultured. (B-C) The development of the arachnid parasite *Tarantobelus* can be robustly sustained using this protocol. (B) *Tarantobelus* adults from the fifth day of axenic culture. Scale bar: 200 µm. (C) The lengths of *Tarantobelus* were measured each day after embryos were deposited in the media. Values are mean ± SD (n= 24 – 25 worms measured for each time point).

It is likely that the shared ability of parasitic nematodes to thrive on simple axenic media reflects adaptations in their feeding mechanisms that are important to their survival in their animal hosts. Free-living nematodes such as *C. elegans* have been described as filter feeders that draw in and eject liquid while trapping particles such as bacteria,^45,46^ and may therefore be poorly equipped to absorb liquid-based nutrients.^47^

### Axenic *S. hermaphroditum* infective juveniles can infect, grow, and reproduce in living insects

We tested our axenically grown *S. hermaphroditum* IJs for their ability to infect and kill insects, using the waxmoth *Galleria mellonella*. When IJs bearing their *Xenorhabdus* bacterial symbiont infect waxmoth larvae, they efficiently kill the insects and rapidly grow to become large reproductive adults (Figure 5A). As previously reported for infection with low numbers of *Steinernema carpocapsae* infective juveniles lacking their pathogenic bacterial symbiont,^48^ axenic *S. hermaphroditum* IJs were not efficient killers of *Galleria* waxmoth larvae: administration of six axenic IJs did not cause an increase in the incidence or speed of the waxmoth larvae’s deaths (10% lethality in six days; n = 20). We then examined the surviving waxmoth larvae to determine what happened to these entomopathogenic nematodes when they had been deprived of their pathogenic capacity, finding that they grew to reproductive adulthood within the living insects (Figure 5B and Video S1). *S. hermaphroditum* populations in living insects could produce a large number of viable, rapidly growing progeny in the time before the larva pupated (Figure 5C). Administration of a single axenic IJ would sometimes (22%; n = 9) produce a colony of *Steinernema hermaphroditum* inside a living insect; at higher doses, infection was more consistently reproducible (adult *S. hermaphroditum* were found in 75% of insects exposed to 1-2 dozen axenic IJs; n = 24). The F_1_ progeny of the adults that had invaded as IJs grew rapidly through larval development within living waxmoth larvae.

**Figure 5.**
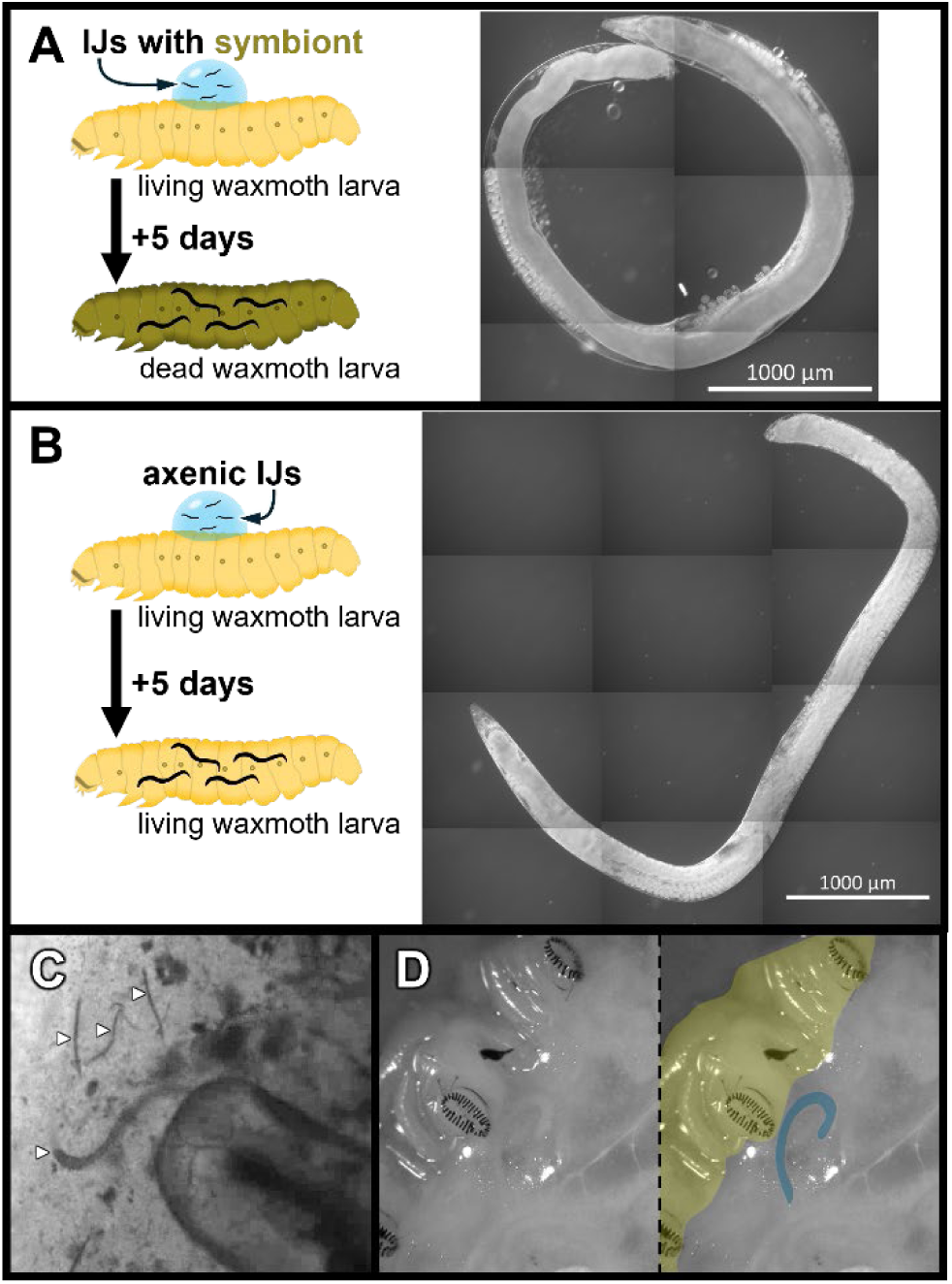
Axenic *S. hermaphroditum* infective juveniles can infect, grow, and reproduce in living insects. (A) Illustration of infection of an insect by an entomopathogenic nematode and its symbiont: *S. hermaphroditum* IJs carrying their *Xenorhabdus* bacterial symbiont were placed on a living *Galleria mellonella* waxmoth larva. Upon entering the insect, they released their symbiont, which rapidly killed and digested the insect. Five days later, adult worms were removed from the carcass; one is shown in the photomicrograph at right with a 1 mm scale bar; the worm is approximately 6.56 mm long. (B) Illustration of infection with axenic IJs. Five days after the IJs were placed on the insect, the insect remains alive and healthy, but within it, adult nematodes and their progeny are found, as shown in the photomicrograph at right, with a 1 mm scale bar; the worm is approximately 6.42 mm long. C) A mixed-stage population of *S. hermaphroditum* animals spilling from a *Galleria mellonella* waxmoth larva, 11 days after infection with IJs, immediately after the living insect was killed. Worms are indicated with white arrowheads. D) An adult worm can be seen in the hemolymph immediately below the cuticle of a partially dissected insect larva, four days after administration of IJs, in a still from Supplementary Video 2. The same photomicrograph is shown twice; on the right, important features are highlighted with colors: the worm is indicated in blue, and the ventral cuticle, including three foot pads, is indicated in yellow. Intact tissues can be seen below the worm.

Axenically raised *S. hermaphroditum* IJs recovered and grew within living waxmoth larvae at a rate comparable to what was seen when waxmoth larvae were treated with, and rapidly killed by, *S. hermaphroditum* larvae that had been grown on their symbiotic *Xenorhabdus* bacteria, and the axenically produced nematodes grew to a size comparable to that seen for infection with symbiotic IJs (∼6.5 mm by day five; Figures 5A and 5B). Although the body lengths of representative *S. hermaphroditum* recovered from insects infected with axenic or symbiotic IJs were similar, there were differences in their appearance: the nematodes growing with the assistance of their bacterial symbiont were fatter and contained more embryos, but also were less robust and more easily damaged when removed from the insect. Dissection of living *Galleria* larvae that contained growing *S. hermaphroditum* showed that the nematodes were present in the fatty tissues directly beneath the cuticle (Figure 5D and Video S2). Inspection using Nomarski differential interference microscopy failed to detect bacteria in the intestines of adult nematodes removed from living *Galleria* larvae following infection with axenic IJs; by comparison, the intestines of nematodes removed from the carcasses of *Galleria* larvae killed by the nematodes’ pathogenic *Xenorhabdus* symbiont contained large amounts of bacteria readily visible using Nomarski microscopy (Figure S2).

The ability of axenic *Steinernema hermaphroditum* IJs to recover and grow to adulthood in living insect larvae was not limited to one insect species, *Galleria mellonella*, but was also seen in *Bombyx mori* silkmoth larvae. Unlike waxmoth larvae, silkmoth larvae were reliably killed by infection with axenic *S. hermaphroditum* IJs: 91% were dead four days after the administration of IJs (n = 56), as high a level of lethality as was seen with *S. hermaphroditum* IJs carrying their pathogenic *Xenorhabdus* symbiont (83% dead after four days; n = 24); these compare to 16% of control larvae that died in the same period (n = 16). Examination of the carcasses of silkmoth larvae that died following administration of axenic IJs showed that 48/51 contained adult nematodes, while the 5 silkmoth larvae that had survived exposure to IJs did not contain nematodes. Because we recovered adult nematodes on the fourth day from the carcasses of silkmoths that, in many cases, had been alive on the third day, those nematodes must have been growing and developing to adulthood prior to the deaths of the infected silkmoth larvae. Unexpectedly, the *Xenorhabdus*-killed silkmoth larvae were an inferior environment for the growth of *S. hermaphroditum*: only 10 of these 20 silkmoth carcasses contained colonies of nematodes. Other insects were not successfully infected by axenic *S. hermaphroditum* IJs: we saw no infection of *Tenebrio molitor* mealworm beetle larvae (n = 3) or *Manduca Sexta* hornmoth larvae (n = 9 unfed and n = 12 well fed); administration of IJs and their *Xenorhabdus* symbiont rapidly killed both *T. molitor* and *M. sexta* insect larvae, and nematode cultures were found in the carcasses.

Having found that *S. hermaphroditum* could infect and grow in living insects, we tested whether other entomopathogens within the same genus shared this capacity. We cultured the dioecious species *Steinernema carpocapsae*, *feltiae*, and *glaseri* to obtain asymbiotic IJs lacking their *Xenorhabdus* symbionts, and used the IJs to infect *Galleria* waxmoth larvae. We found that nematodes of all three species developed to adulthood within living waxmoth larvae (Figure S3) and that all three species were able to mate and reproduce within living insects.

### *H. bacteriophora* can infect and grow in living insects independent of their symbiotic bacteria

We also tested *Heterorhabditis bacteriophora*, representing the other genus that consists entirely of entomopathogens, using symbiont-free IJs grown on a colonization-defective mutant strain of *Photorhabdus temperata* (see Methods). We found that symbiont-free *Heterorhabditis* nematodes developed to adults inside living *Galleria* larvae, indicating they were capable of feeding inside living insects in the absence of their symbiont, but that these adults were dramatically smaller than the nematodes recovered from *Galleria* larvae that had been manually infected with *Photorhabdus* (Figure 6). This is consistent with earlier reports that *Heterorhabditis* lacking their symbionts can grow to fertile adults in *Galleria* larvae.^49^

**Figure 6.**
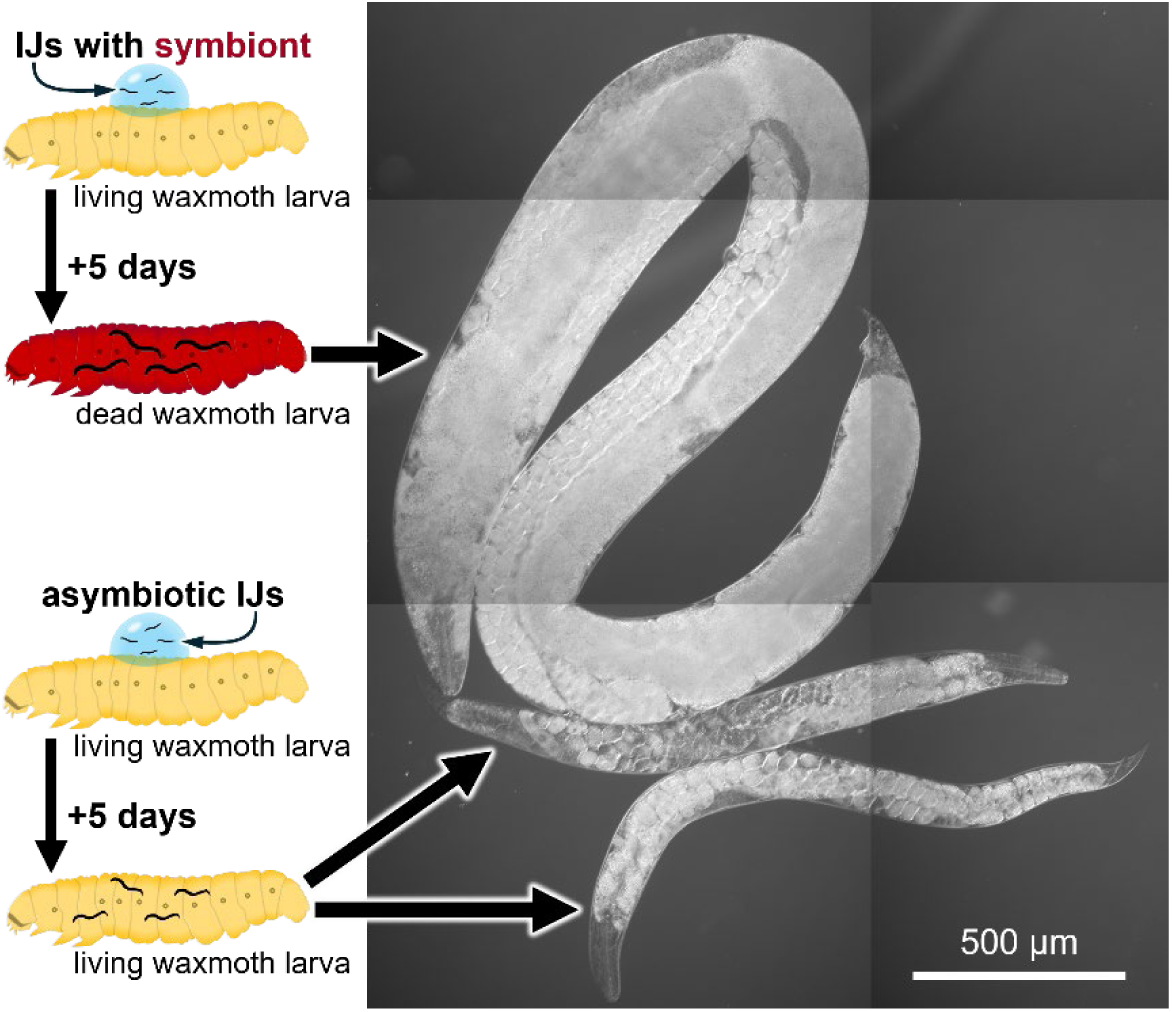
*Heterorhabditis bacteriophora* adults removed from *Galleria mellonella* waxmoth larvae five days after infection with IJs. The large adult at the top developed from an IJ that infected an insect treated with, and subsequently killed and digested by, its *Photorhabdus* bacterial symbiont. The two much smaller adults below were recovered from a living *Galleria* larva infected with asymbiotic IJs. They can be seen to contain embryos, some of which are in the four-cell stage.

The *H. bacteriophora* adults recovered from these living *Galleria* larvae contained embryos that appeared upon inspection to be undergoing normal early development, but no viable F_1_ progeny were found when *H. bacteriophora* adults were seen in living *Galleria* that had been infected with asymbiotic IJs in repeated trials (n > 70 adults without viable progeny). When adult *H. bacteriophora* were recovered from living *Galleria* larvae and transferred to lawns of *Photorhabdus* bacteria, they were able to produce viable progeny, starting after two days of feeding on *Photorhabdus*. These observations are consistent with previous reports that *Heterorhabditis* require unknown substances from their *Photorhabdus* symbiont for development. The ability of *H. bacteriophora* and multiple species of *Steinernema* to grow to fertile adulthood solely by feeding on a living insect host indicates that, before they became committed to feeding on their bacterial symbionts in an entomopathogenic lifestyle, both genera were adapted to feeding on the nutrients of living insects as parasites or parasitoids.

### *Steinernema* can emerge from insect hosts carrying bacteria acquired in the infection

We tested *Steinernema hermaphroditum* IJs recovered from insect carcasses for colonization by bacteria. We collected IJs from the carcasses of insects that had been infected either with symbiotic or axenic IJs, washed the IJs in a bleach solution to remove bacteria from their surface, and examined them for bacteria by using PCR to amplify bacterial 16S ribosomal DNA and by culturing their bacteria on LB media (Fig. 7 and see Methods). As expected, symbiotic infection resulted in IJs carrying viable colonies of *Xenorhabdus griffiniae*. We found that two different waxmoth infections with axenic IJs produced IJ populations carrying identical or closely related isolates of *Serratia marcescens*. Tests of IJs produced from silkmoths resulted in three sequences, two of them identical, closely matching *Serratia marcescens* and one matching *Shouchella clausii*. We found that the waxmoth-derived strain of *Serratia marcescens* was not a suitable food source for *Steinernema* (Figure S4) and did not confer insect-killing ability upon the IJs.

**Figure 7.**
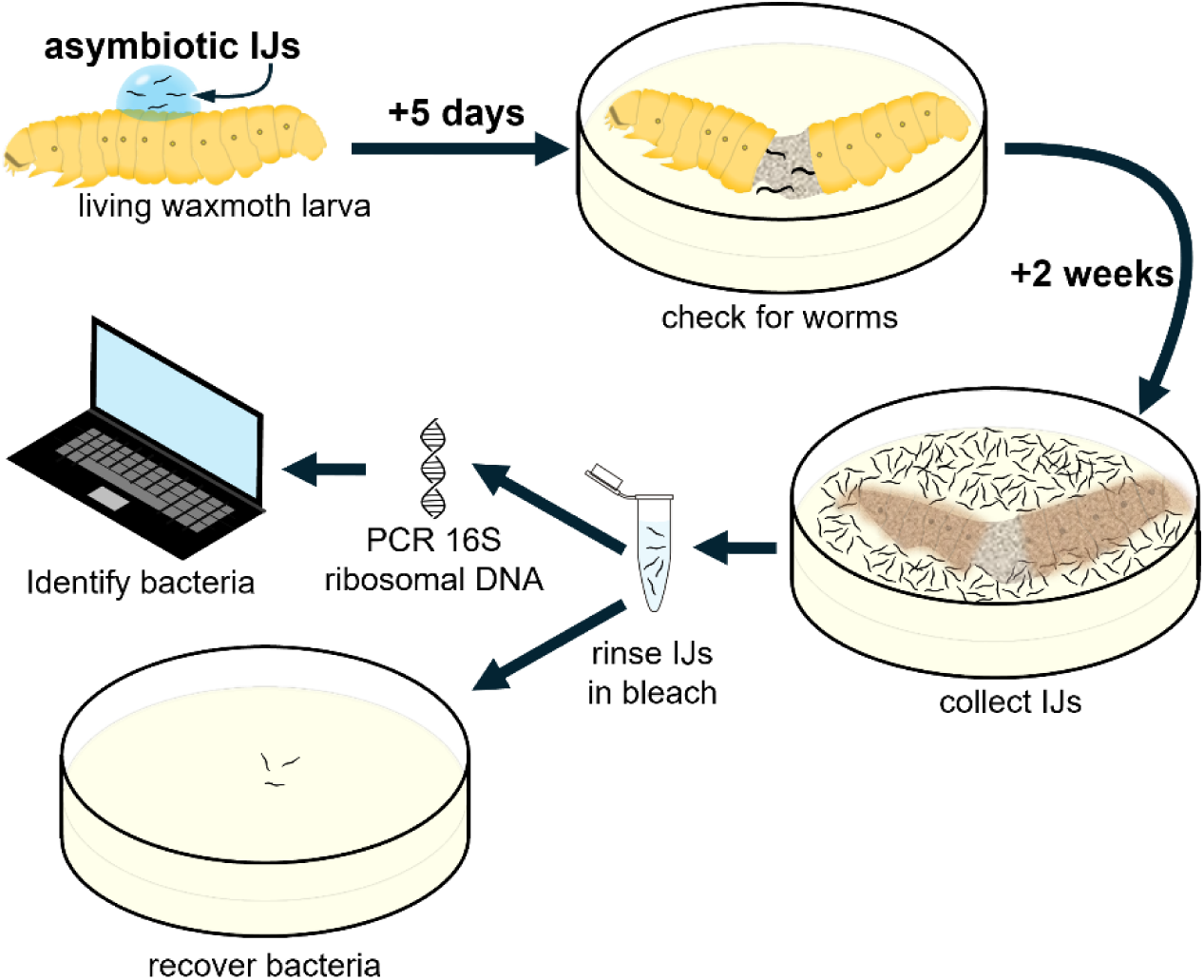
Identification of bacteria associated with IJs recovered after infection of waxmoth larvae with axenic IJs. After insects are sacrificed to confirm they are colonized by nematodes, the carcasses and their worms are incubated for two weeks to allow the nematode population to produce IJs. IJs are recovered, and their surface bacteria cleaned using a bleach solution, washed, and then either processed for bacterial 16S ribosomal DNA sequence or cultured to recover their bacteria.

## Discussion

In this study, we explored the evolutionary adaptation of parasites, focusing on two species: *Steinernema hermaphroditum* and *Heterorhabditis bacteriophora*, representatives of two independent groups of entomopathogenic nematodes that have converged on a highly distinctive lifestyle despite being separated by hundreds of millions of years. To study this, we developed an axenic culture medium that allows for robust and sustained growth of *S. hermaphroditum* and the production of its infective juvenile (IJ) larvae (Figure 1). We subsequently found that the ability of nematodes to utilize this culture medium appeared to be common among the parasitic nematodes we tested, but was largely absent in the free-living nematodes we tested. It therefore seems that as nematodes develop the capability to make use of animal hosts as parasites, they undergo adaptations in their feeding that also make them culturable using this simple growth medium. Parallels can be drawn between this and plant-parasitic nematodes that have repeatedly and independently evolved a stylet capable of puncturing plant tissues and enabling them to feed upon their host.^7^

As parasites continue to evolve, they can develop a life cycle that never requires them to grow outside of their preferred single type of host organism and thereby become obligate parasites. Traits that were advantageous when they lived in the outside world or in a complex-lifecycle parasite whose life cycle involves a progression of hosts will not be selectively maintained and can easily be lost.^50^ The loss of traits that are no longer needed can even be reproducible or predictable, as in the examples of lipid synthesis capabilities and egg yolk genes that have repeatedly been lost in the independent evolution of multiple species of parasitoid wasp.^1,51^ We found an example of this in the inability of our entomopathogenic nematodes to survive hatching in the absence of food. This stands in contrast to the well-studied free-living saprophytic nematode *Caenorhabditis elegans* and its close relatives, which are known to developmentally arrest when they hatch in the absence of food and to seek out food sources.^30–33,52^ We found that these abilities are broadly conserved across a wide survey of free-living nematodes, but that both *Steinernema* and *Heterorhabditis* nematodes had lost this capacity.

The embryos of *Steinernema* and *Heterorhabditis* only ever hatch inside of a host, in the presence of ample resources, and more specifically in the presence of a near-monoculture of the bacterial symbiont that has evolved to serve as their food source and will later enable them or their children to kill and efficiently consume an insect. If they ever hatched outside of this environment, they could not hope to explore and find their way into a new host, a role that is performed by specialized IJs later in larval development. This could explain why there is no evolutionary pressure to maintain survival capability in food-deprived hatchlings, whereas this is observed in every other nematode genus we tested. Not only was this capability lost in both *Steinernema* and *Heterorhabditis* nematodes as they evolved as obligate parasites, but in each genus it was lost differently: *Steinernema* hatchlings could not go into developmental arrest and rapidly starved to death, but retained some remnant ability to disperse in search of food sources, albeit barely, while *Heterorhabditis* nematodes could still arrest their development and recover when food was provided – albeit much worse than any free-living nematode – but their hatchlings completely lack food-seeking behavior. The same convergent evolution that caused *Steinernema* and *Heterorhabditis* nematodes to acquire similar traits, such as the ability to grow by feeding on hemolymph or in our simplified axenic media, also caused both entomopathogens to independently and differently develop the capacity for hatchlings to overcome the absence of food, made unnecessary by their obligate-parasitic lifestyle.

On the other hand, to exploit the rich resource environment of the interior of an insect host, the entomopathogenic nematodes developed the capacity to feed independently of bacteria, possibly through adaptations that also allowed them to grow in our simple axenic media. *Steinernema,* in particular, grows as rapidly and to a similarly large size when growing in a living insect as it does when feeding on its bacterial symbiont in the carcass of an insect their bacteria symbiont killed and digested for them. Dissection shows that the worms grow in the insect’s hemolymph, where they are likely feeding directly on the insect, perhaps drinking the hemolymph much as they drink our modified bacterial growth media. When grown in the hemolymph there are no bacteria visible in the nematode, and any persistent bacterial infection in the hemolymph would likely have killed the apparently healthy insect.

The ability of entomopathogenic nematodes to invade a living insect and feed directly on hemolymph without their associated symbiotic bacteria suggests their distant ancestors were parasites or parasitoids of living insects (see Graphical Abstract), which has been an area of speculation. ^19^ We found that worms that entered insects free of bacteria could later produce dispersing IJs that carried bacteria they found within. This included *Serratia marcesens*, strains of which have been reported to be associated with multiple genera of entomopathogenic or occasionally entomopathogenic nematodes^53–55^ and can function as a pathogen of insects.^56^ The particular strain of *Serratia* we found was not a suitable food source for our worms and did not seem to enhance their ability to kill insects, but at some point in the distant past a common ancestor of all *Steinernema* must have been more fortunate in the bacteria it carried out of an insect, and began a partnership with *Xenorhabdus* bacteria that has been found in all 114 *Steinernema* species so far described.^57,58^ We suspect *Heterorhabditis* has a similar history, somewhat obscured by the extremely close partnership between *Heterorhabditis* and its bacterial symbiont *Photorhabdus*: at some point, *Heterorhabditis* became completely dependent on *Photorhabdus* for development,^42–44^ which limited its ability to grow and reproduce both in our axenic culture and in living insects. It may be possible to determine the nature of this dependence by experimenting with supplements to our culture medium.

Overall, our systematic analysis of the evolutionary adaptation of entomopathogenic nematodes reveals striking examples of convergent evolution both traits gained (feeding capacities shared among widely varying parasitic nematodes) and lost (both of our obligate parasites independently lost the ability of hatchlings to overcome food deprivation) and provides strong evidence for a parasitoid origin of entomopathogenic nematodes and a mechanism for their later partnership with pathogenic bacteria.

## Acknowledgments

We thank the following for providing nematode and bacterial strains: Anil Baniya and Adler Dillman (UC Riverside) and Pierluigi Perfetto, Eustachio Tarasco, and Daniele Cornara (U of Bari Aldo Moro) for *Pellioditis hermaphrodita*, *Steinernema carpocapsae*, *Steinernema feltiae*, *Steinernema glaseri*, and *Tarantobelus jeffdanielsi*; James Lee and Cori Bargmann (Rockefeller U) for *Auanema freiburgensis*; and Ryoji Shinya (Meiji U) for *Bursaphelenchus okinawaensis*. We thank Joseph Parker (Caltech) for instruction in dissecting insects; Jennifer Heppert (U Tennessee), Adler Dillman, and Erich Schwarz (Cornell) for valuable suggestions; Yanping Qin and Tsui-Fen Chou (Caltech) for technical assistance. Some strains were provided by the CGC, which is funded by the NIH Office of Research Infrastructure Programs (P40 OD010440). This work was supported by NSF EDGE grant 2128267 (to PWS) and Caltech’s Center for Evolutionary Science (CES) and Center for Environmental Microbial Interactions (CEMI). This work was also facilitated by WormBase, a knowledgebase for nematode research ^59^, and Nemys: World Database of Nematodes ^60^.

## Methods

### Key resources table

| REAGENT or RESOURCE | SOURCE | IDENTIFIER |
| --- | --- | --- |
| **Bacterial strains** |  |  |
| *Comamonas aquatica* | Caenorhabditis Genetics Center | DA1877<sup>46</sup> |
| *Escherichia coli* | Caenorhabditis Genetics Center | OP50<sup>61</sup> |
| *Xenorhabdus griffinae* | Sternberg Lab | HGB2511<sup>22</sup> |
| *Photorhabdus temperata* | Sternberg Lab | NC1-TRN16<sup>62</sup> |
| **Chemicals** |  |  |
| Hemin | Sigma-Aldrich | H9039 |
| **Critical commercial assays** |  |  |
| QiaQuick PCR purification kits | Qiagen |  |
| **Deposited data** |  |  |
| *X. griffinae* 16S | This study | PX425005 |
| Waxmoth *Serratia* 1 | This study | PX425006 |
| Waxmoth *Serratia* 2 | This study | PX425007 |
| Silkmoth *Serratia* 1 | This study | PX425008 |
| Silkmoth *Serratia* 2 | This study | PX425009 |
| Silkmoth *Serratia* 3 | This study | PX425010 |
| Silkmoth *Shouchella* | This study | PX425011 |
| **Experimental models:**<br>**Organisms/strains** |  |  |
| *Steinernema hermaphroditum*<br>inbred wild-type strain | Sternberg Lab | PS9179<sup>22</sup> |
| *Steinernema felitae* wild-type strain | Adler Dillman | SN<sup>63</sup> |
| *Steinernema glaseri* wild-type<br>strain | Adler Dillman | NC<sup>63</sup> |
| *Steinernema carpocapsae* (Italy)<br>wild-type strain | Adler Dillman | APU-1 |
| *Steinernema carpocapsae* (Poland)<br>wild-type strain | Adler Dillman | PLB-18 |
| *Bursaphelenchus okinawaensis*<br>wild-type strain | Ryoji Shinya | SH1<sup>64</sup> |
| *Acrobeloides buschlii*<br>wild-type strain | Caenorhabditis Genetics Center | DWF1107<sup>65</sup> |
| *Panagrolaimus* sp. PS1159<br>wild-type strain | Sternberg Lab | PS1159<sup>66</sup> |
| *Tarantobelus jeffdanielsi* wild-type<br>strain<sup>37</sup> | Adler Dillman |  |
| *Caenorhabditis elegans*<br>wild-type strain | Caenorhabditis Genetics Center | N2<sup>61</sup> |
| *Pellioditis hermaphrodita* wild-type<br>strain | Adler Dillman | ITD272<sup>67</sup> |
| *Pellioditis pelhamensis* | Caenorhabditis Genetics Center | EM434 |
| *Auanema freiburgensis* wild-type<br>strain | James Lee /Cornelia Bargmann | SB372<sup>68</sup> |
| *Heterorhabditis bacteriophora*<br>inbred wild-type strain | Sternberg Lab | M31e<sup>69</sup> |
| *Oscheius tipulae* wild-type strain | Caenorhabditis Genetics Center | CEW1<sup>70</sup> |
| *Pristionchus pacificus* wild-type<br>strain | Caenorhabditis Genetics Center | PS312<sup>71</sup> |
| *Saccharomyces cerevisiae* | Sternberg Lab | W303-1A |
| *Galleria mellonella*<br>waxmoth larvae | PetSmart (Pasadena, CA) |  |
| *Tenebrio molitor*<br>mealworm beetle larvae | PetSmart (Pasadena, CA) |  |
| *Manduca sexta*<br>hornmoth larvae | Reptile Supply (Wichita, KS) |  |
| *Bombyx mori*<br>silkworm larvae | The Silkworm Shop (San Diego,<br>CA) |  |
| **Software and algorithms** |  |  |
| Fiji (ImageJ)<sup>72,73</sup> | National Institutes of Health |  |

### Nematode culture (bacterial-based culture)

*Steinernema* strains were grown in Petri plates at 25°C using the *Comamonas aquatica* strain DA1877^46^ on Enriched Peptone Media (EPM) agar as described,^24^ or *Xenorhabdus griffiniae* strain HGB2511^22^ on Nematode Growth Media (NGM) agar as described.^22^ *Bursaphelenchus okinawensis* was grown at 25°C using *Saccharomyces cerevisiae* strain W303-1A^74^ as a food source on Malt Extract Agar media as described.^64^ *Heterorhabditis bacteriophora* was grown at 25°C using *Photorhabdus temperata* strain NC1-TRN16^62^ as a food source on Lipid Agar media ^75^. All other strains were grown at 20°C or at room temperature using *Escherichia coli* strain OP50 as a food source on NGM agar as described.^61^ Bacterial-based (*C. aquatica* DA1877) liquid culture was done similarly to what has been described.^26^ Briefly, overnight cultures of DA1877, seeded by overnight cultures in a smaller volume, were pelleted through centrifugation and re-suspended in S medium, 1/10 of the culture volume.

### Preparation of axenic, nutrient-free nematode eggs

Strains were prepared for attempts at axenic culture by washing eggs and gravid nematodes from Petri plates in water, centrifuging to pellet the eggs and mothers, rinsing well in water, and using bleach solutions to clean the eggs and dissolve the gravid mothers: either 0.8 M NaOH with 5% sodium hypochlorite solution or 0.5 M NaOH with ∼8% sodium hypochlorite solution (Household bleach solution). Eggs and mothers in bleach solution were agitated until the mothers had visibly dissolved, which sometimes included pelleting the eggs by centrifugation to aspirate and replace the bleach solution. After the bleach treatment was complete, the eggs were rinsed either in sterile water or M9 buffer (For 1 liter: 3 g KH_2_PO_4_, 6 g Na_2_HPO_4_, 5 g NaCl, and with 1 ml 1 M MgSO_4_ added after autoclaving) at least twice, after using centrifugation to pellet the eggs and removing the supernatant by aspiration.

### Preparing the axenic media

The axenic culture media, with composition as described in Figure 1A, were prepared by mixing the following 6 pre-made solutions and water:

(1) Carbon source, consisting of tryptone (100 g/L) and yeast extract (50 g/L) in the same proportion as LB (essentially, a 10XLB solution without NaCl). Autoclaved to sterilize and stored at room temperature. (2) NaCl solution, consisting of NaCl (50 g/L) (10X the normal salt solution in LB). Autoclaved to sterilize and stored at room temperature. (3) Trace metal solution adapted from *C. elegans* protocols as described in Stiernagle ^28^, consisting of: FeSO_4_·7H_2_O (0.69 g/L), MnCl_2_·4H_2_O (0.2 g/L), ZnSO_4_·7H_2_O (0.29 g/L), CuSO_4_·5H_2_O (0.025 g/L), and Disodium EDTA (1.86 g/L). Autoclaved to sterilize and stored in the dark at room temperature. (4) Cholesterol solution, from *C. elegans* protocols as described in Stiernagle ^28^: cholesterol (5 mg/mL) in ethanol. Filter sterilized and stored at room temperature. (5) Hemin solution: hemin (Sigma, H9039) 10mM in DMSO. Stored at −20°C, precipitation was often observed when thawed. (6) Carbenicillin solution (25 mg/mL). Filter sterilized and stored at −20°C.

For 10 mL of the axenic media (Growth): add 3.2 mL of the carbon source solution, 1 mL of the NaCl solution (1:10), 0.1 mL of the trace metal solution (1:100), 0.01 mL of the cholesterol solution (1:1000), 0.02 mL of the hemin solution (1:500), 0.02 mL of the carbenicillin solution (1:500), and add water to 10 mL (in this case 5.65 mL). The amount of carbon source solution can be adjusted to suit the needs of the culture. In “Infective Juvenile Inducing” media (Figures 1F and 1G), the carbon source solution is reduced to 1 mL, while the other ingredients remain constant, with water to 10 mL. In the nutrient reduction experiments (Figure 2A), the carbon source solution was reduced progressively from 1 mL (100% nutrient) to 0 mL (0% nutrient). In the experiments described in Figure 4, three versions of the media were tested for all species, with carbon source media at concentrations of 3.2 mL, 1.6 mL, and 1.0 mL per 10 mL of media. All three parasitic species that could be readily cultured with the media grew best in media prepared containing 3.2 mL of carbon source media per 10 mL.

### Liquid culture of nematodes

Eggs axenized as above were transferred to the axenic culture media or the bacterial-based liquid culture and incubated either at 25°C or 20°C in 6 to 24-well plates or Erlenmeyer flasks. In all cases, the culture was maintained with shaking at 60 rpm, but this may not be necessary. When using tissue culture plates, the plates were incubated in humidified chambers made by placing wet paper towels in plastic shoe boxes, similar to what we previously described. ^26^ The paper towels were wetted regularly to maintain humidity. All the experiments described in Figures 1 C-G were conducted at 25°C. In Figure 4A, the first rounds of analysis for *A. buschlii, Panagrolaimus sp. PS1159, T. jeffdanielsi, C. elegans, P. hermaphrodita, A. freiburgensis, O. tipulae,* and *P. pacificus* were conducted at 20°C, and *S. hermaphroditum*, *B. okinawaensis*, and *H. bacteriophora* were conducted at 25°C. Of the worms that were incubated at 20°C, all but *A. buschlii* were assayed again at 25°C (including data for Figures 4B and 4C), as we confirmed that all the worms in our panel can be cultured at the elevated temperature (Figure S5).

Imaging of the worms was done by dropping a small volume of the worm-containing culture medium onto 2% agarose pads on microscopic slides. The mounted worms were then observed using a Zeiss Imager Z2 microscope, and the images (Figures 1D, F, 4B, S1B-C) were captured through an Axiocam 506 mono and Zen 2 Blue software. To measure the growth of the worm (Figures 1C, 4C, S1A, S1C), worm lengths were measured by carefully drawing a segmented line from head to tail through the midline of each worm using ImageJ (NIH) software as previously described^22,76^. At stages when the sex of the nematode could be determined, only those with the hermaphroditic/female body type were measured. In Figures 1F and 1G, IJ were determined by the enclosure of the worm in a detached cuticle.

### Nematode hatchlings arrest and recovery assay

#### Experimental setup

Axenic eggs of various nematode species were allowed to hatch in M9 buffer, incubated at 25°C or 20°C. When we were uncertain whether a species could grow well and produce healthy progeny at 25°C, we placed eggs on lawns of OP50 bacteria on NGM agar and observed the populations’ development once per day for seven days (Figure S5). With the exception of *B. okinawaensis* (see below), worms containing M9 buffer were transferred to 15-mL conical tubes (3 mL/tube). Each species is assayed in 4 biological replicates (4 tubes). At each time point, aliquots of the culture were taken out for survival and recovery assessments. In all assays, if multiple trials were performed for the same species, the trial with the highest number of assayed animals was analyzed.

As the experiment is to assess the hatchling’s ability to endure without food, we took caution to prevent the presence of significant worm debris in the culture, as it could be utilized as a food source for some nematodes. This is very difficult in some of the species, while impossible with *B. okinawaensis*. Therefore, for *B. okinawaensis*, embryos and newly hatched larvae were recovered using glass micropipettes, and the survival assessment was conducted differently (see below). In addition, because young larvae of *B. okinawaensis* could not be reliably seen in the opaque yeast lawns used as their food source, they were not included in the recovery assay.

#### Survival assessment

To assess worm survival, small drops of the culture were placed on agar plates (Chemotaxis or NGM) without bacterial lawns, and the movement and morphology of the worms within the liquid droplets were assayed using a dissecting microscope. The survival of the worms was predominantly assessed based on their movement, assisted by their morphology. The worm mobility in Figure 2A was also assessed similarly using this method. In some cases, worms could be scored alive if their general morphology resembles that of mobile worms. In a few species (*A. buschlii*, *T. jeffdanielsi, O. tipulae*), the movement of a large percentage of the worms ceased or decreased dramatically a few days into the experiments, but they had otherwise morphological characteristics that resembled living worms. In such cases, suspected worms were isolated and allowed to recover on food. If most of such worms were able to recover (See note 1 below), they were undoubtedly alive, and other worms with similar morphological characteristics in that species would be scored as so, despite the general lack of movement. In some of the species, some of the hatchlings could shrink and have a rugged appearance, but they would still be scored as alive if movement could be observed, or in the case of *O. tipulae* (Note 2), scored as alive even if no movement could be observed. 20 – 325 worms were scored for each data point (each tube at each time point). Carcasses of larvae that appeared to have hatched and died during the axenic egg preparation process were not included in the assay.

Note 1: *A. buschlii* that had appeared completely motionless in liquid media recovered after being given food, in the following proportions, transferred the indicated number of days after the eggs were prepared: 31/35 (day 4) and 34/36 (day 9). Similarly, motionless *T. jeffdanielsi* recovered after being given food in the following proportions and on the indicated days: 24/26 (day 2), 17/20 (day 6), 21/25 (day 12). Similarly motionless (but otherwise normal appearing) *O. tipulae* recovered after being given food in the following proportions and on the indicated days: 25/26 (day 1), 25/26 (day 4), and 25/25 (day 9).

Note 2: 24 of 25 *O. tipulae* that were motionless and had adopted a shrunken and ridged appearance after nine days of food deprivation from hatching recovered after being given food.

For *B. okinawaensis*, embryos and newly hatched larvae were recovered from axenic egg preparation using glass micropipettes to separate them from worm debris that was not completely dissolved by bleaching. The slow manual isolation prevented us from performing exactly the same survival assessment as was used for other species, in which each worm was only ever scored once. *B. okinawaensis* hatchlings were scored repeatedly at different time points by observing the worms in 96 to 12 well tissue culture dishes. Worms were transferred between wells and dishes to assist the assay. We did not score all the worms of each of the 4 groups (replicates) at each time point, as finding a small and thin moving worm in liquid was very difficult (which is why other species were assayed on plates), and this is likely the reason for the variation near the end of the *Bursaphelenchus* experiments.

#### Recovery assessment

To assess the recovery of the worms, aliquots of the culture were taken out at each time point and seeded onto plates seeded with food corresponding to the diet of the species. 3 plates were seeded per tube, and 12 plates (3 plates x 4 tubes) were seeded per species per time point. Worms were given time to develop, and the number of worms on each plate was counted. Counted worms were removed to avoid double-counting. When possible, after removing all counted worms, we assessed the plates daily for worms that may have delayed recoveries, which we observed in *C. elegans* and *P. hermaphrodita* after prolonged starvation. The number of worms from the 3 plates of each tube was then averaged, with the resulting number used as a single data point representing the tube (biological replicate) at that specific time point. The numbers corrected for the volume of the culture seeded were then compared to the numbers obtained at day 0, which was seeded immediately at the start of the experiment. In some cases - *A. buschlii, A. freiburgensis* (20°C and 25°C), *P. hermaphrodita* (20°C), and *C. elegans* (20°C), many more worms were found in the first data points compared to that of the time 0 control. This is likely due to the fact that at day zero, immediately after the egg preparation, most of the animals in the media are embryos, which likely clump and attach to the walls of the tube differently compared with the rest of the experiments, for which the animals largely consist of hatchlings. To correct for this, we estimated the number of expected animals at day 0 using the number of animals in the closest time point and the survival percentage we obtained through the survival assessment assay described above, when such incidences occurred. At least 20 worms total were scored from the three control plates of each tube; in most cases, > 20 worms were scored from each individual plate. Overall, across all species and temperature combinations, 97-3,145 worms were counted at time 0 for the controls in each experimental set. The long-term endurance experiments were performed without the use of antibiotics, as the media contains no nutrients and does not allow microbial growth; however, sometimes contamination can be seen in the recovery assays, in which the media was spotted onto rich agar media, and data points were abandoned in cases where the contamination is likely to significantly impact the survival of the worm or obstruct observations. As a result, a few experiments contained timepoints in which there were only 3 biological replicates.

In both the survival and recovery assessments, we attempted to score the worm as close to the precise timepoints as possible, particularly at the start of the assay. A major exception is *A. buschlii* at the day 30 timepoint, which was assayed 13 hours late.

### Food-seeking assay

8.5 cm Petri plates were prepared containing chemotaxis agar.^77^ *Comamonas* DA1877 bacteria were grown overnight in LB, centrifuged, and resuspended in water at 1/10 of the original culture volume. *Photorhabdus* NC1-TRN16 bacteria were scraped in a small volume of M9 buffer from heavily pigmented lawns grown on lipid agar and centrifuged, and the pellet was resuspended in an equal volume of M9 buffer. 15-20 µl of bacterial suspension was deposited on an 8.5 cm chemotaxis agar Petri plate to create a lawn approximately 0.5-0.7 cm in diameter, 2 cm from the edge of the plate along a line that bisected the plate (Figure 3A). Control populations of axenized eggs were placed either directly on the lawn of a food-seeking assay plate (*H. bacteriophora*), on lawns of DA1877 on EPM in Petri plates (*S. hermaphroditum*), on lawns of OP50 on NGM in Petri plates (all other species), and the number of animals that grew from these eggs was counted. Experimental populations were placed within a 0.5 cm diameter spot 4 cm away from their food source along the line bisecting the plate, 2 cm from the other edge of the plate. Worms were given 36 and 60 hours at 25°C to reach the bacterial lawn, after which the worms feeding on the lawn, or immediately next to the lawn, were counted. At least four experimental and four control plates were used for each species. Because young larvae of *Bursaphelenchus okinawaensis* could not be reliably seen in the opaque yeast lawns used as their food source, they were not included in this assay.

Axenized eggs were placed either directly on the lawn of the chemotaxis plate (*H.* bacteriophora, as a control to demonstrate their viability and the lawn’s suitability as a food source), on the lawns of DA1877 in EPM plates (*S. hermaphroditum*), on the lawns of OP50 in NGM plates (all other species), or on the chemotaxis plates within a 0.5 cm diameter spot 4.5 cm away along the line bisecting the plate, 2 cm from the other edge of the plate (as experimental plates). Worms were given 36 and 60 hours at 25°C to reach the bacterial lawn, after which the worms feeding in the bacterial lawn were counted. At least four experimental and four control plates were used for each species. Because young larvae of *Bursaphelenchus okinawaensis* could not be reliably seen in the opaque yeast lawns used as their food source, they were not included in this assay.

### Infection of insects

Food-deprived larvae of the waxmoth *Galleria mellonella* and the mealworm beetle *Tenebrio molitor* were purchased at PetSmart (Pasadena, CA) and kept at 15°C until used. Larvae of the hornmoth *Manduca sexta* were purchased from Reptile Supply (Wichita, KS) and used immediately on arrival. Larvae of the silkworm *Bombyx mori* were purchased from The Silkworm Shop (San Diego, CA) and used immediately on arrival. To infect insects, IJs suspended in water were pipetted onto the backs of the insect larvae; control insects had water pipetted onto their backs. In some experiments, insect larvae were photographed daily to monitor their condition. Waxmoth larvae were placed on 5.5 cm Whatman #1 filters in 6 cm Petri plates that had been left undampened or dampened with 200 µl sterile water. Hornmoth larvae were kept as individuals in 6 cm Petri plates containing a 5.5 cm Whatman #1 paper that was regularly moistened with sterile water, or were maintained in the incubators in which they had been shipped: vented plastic containers that had a food source at the top and a climbing frame from the base of the container to the food source. Silkworm larvae were kept in 6 cm Petri plates containing a 5.5 cm Whatman #1 filter or in 6 cm Petri plates not containing a Whatman paper, and were provided with 0.5-1 ml of silkworm food (The Silkworm Shop) that was prepared according to the manufacturer’s instructions and stored at 4°C until used. All incubations of insect larvae with nematodes were done at 25°C. *Steinernema hermaphroditum* IJs used to infect insects were either axenic (isolated from cultures grown in axenic liquid media) or symbiotic (isolated from cultures grown on HGB2511 on NGM agar media), as indicated in the experiments. *Steinernema carpocapsae*, *feltiae*, and *glaseri* IJs used to infect insects were asymbiotic (isolated from cultures grown on the neutral soil bacterium DA1877 on EPM agar media). *Heterorhabditis* IJs were obtained from cultures grown on the non-colonizing *Photorhabdus* strain NC1-TRN16 on Lipid Agar media and were rinsed in 1% commercial bleach to remove any bacteria on their surface, and so were asymbiotic. When it was desirable to reunite axenic or asymbiotic nematodes with their bacterial symbiont within an insect, the insect was stabbed with a 26G needle coated in pathogenic bacteria shortly before IJs were placed upon it; lepidopteran insects (*Bombyx*, *Galleria*, *Manduca*) were stabbed in a foot pad, *Tenebrio* were stabbed through the dorsal cuticle. To look for worms infecting insect larvae, the insects were dismembered in water or in M9 buffer or were dissected by using microdissection scissors to remove the head and cut open the ventral cuticle along the length of the larva. Video recordings were made using a Zeiss M^2^BIO stereomicroscope.

### Isolation and identification of bacteria associated with IJs

One *Galleria* larva manually infected with *X. griffiniae* HGB2511 and infected with axenically grown *S. hermaphroditum* PS9179, and two living *Galleria* larvae infected with axenic PS9179 were dismembered in buffer to confirm they contained colonies of *S. hermaphroditum* and then placed on EPM agar Petri plates and incubated until they produced IJs, which were collected by washing the plate with water. The IJs were pelleted by centrifugation, washed three times in water, and surface-sterilized by exposing them to 1% commercial bleach for 5 minutes, followed by two more washes in sterile water. The collected IJs were split into an aliquot that was lysed for PCR according to a standard *C. elegans* method^32^ and an aliquot that was placed on LB agar plates to recover any bacteria they contained; the bacteria could grow in the carcasses of IJs that did not survive the bleaching and centrifugation, or if an IJ spontaneously emerged from developmental arrest. By colony morphology, each isolate appeared to contain a single type of bacteria that grew well on LB agar at 25°C. PCR was performed on the lysate using *Taq* polymerase and the BacID set of primers^33^; PCR products were purified using QiaQuick (QIAGEN, Germantown, Maryland) and submitted for sequencing (Plasmidsaurus, Redwood City, California). The resulting sequences were identified using NCBI BLAST. The bacterial sequence associated with the worms from the *Xenorhabdus*-killed *Galleria* larva matched *Xenorhabdus griffiniae*, and the bacterial sequences associated with the worms from each of the two living *Galleria* larvae matched *Serratia marcescens*. Additional bacterial sequences were obtained by using BacID primers to sequence bacterial DNA present in lysates of IJs recovered from silkworm larvae killed by infection with axenic *S. hermaphroditum*, after the IJs had been washed in bleach solution; these were identified as three additional isolates of *Serratia marcescens*; two of these were identical, with the third slightly differing in sequence, and both differed from the two isolated from *Galleria*, and one sequence that matched *Shouchella clausii*. The isolate of *Serratia marcescens* was tested for its ability to support *S. hermaphroditum* growth by placing axenized eggs on lawns of the bacteria grown on EPM or NGM and incubating the plates at 25°C, with bare media (NGM without bacteria) and standard growth plates (DA1877 on EPM, and HGB2511 on NGM) as controls. IJs from the population that tested positive for *Serratia marcescens* by PCR and contained viable colonies of *Serratia marcesens* were tested for their ability to kill *Galleria mellonella* larvae, as described above.

---

**Figure S1.**
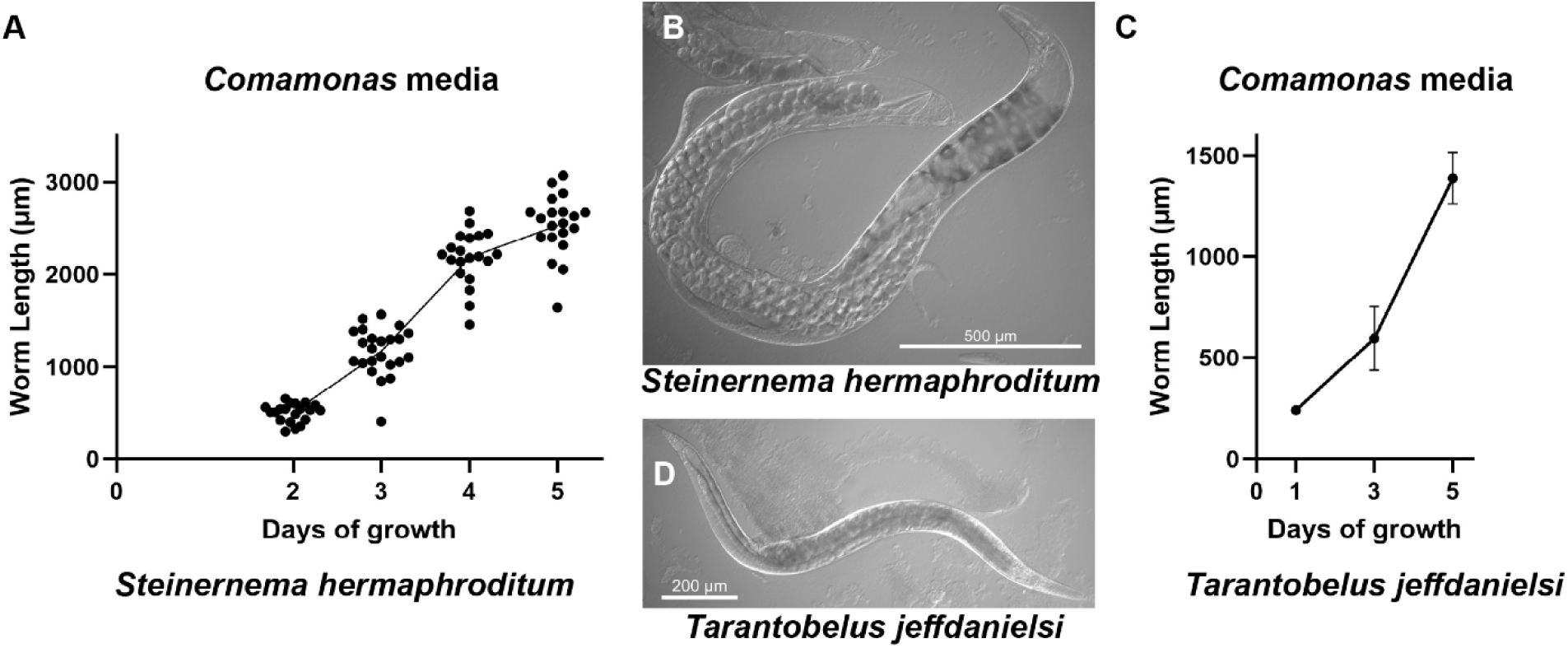
The development of nematodes in bacteria-based liquid culture. Growth of *S. hermaphroditum* (A, B) and *T. jeffdanielsi* (C, D) worms in bacterial-based liquid culture using *Comamonas aquatica* as a food source. The lengths of the worms were measured at each time point after embryos were placed in the culture media. Values are mean ± SD (n= 20 – 31 worms measured for each time point). Photomicrographs (B, D) show adults on the fifth day of growth in culture. Scale bars as indicated.

**Figure S2.**
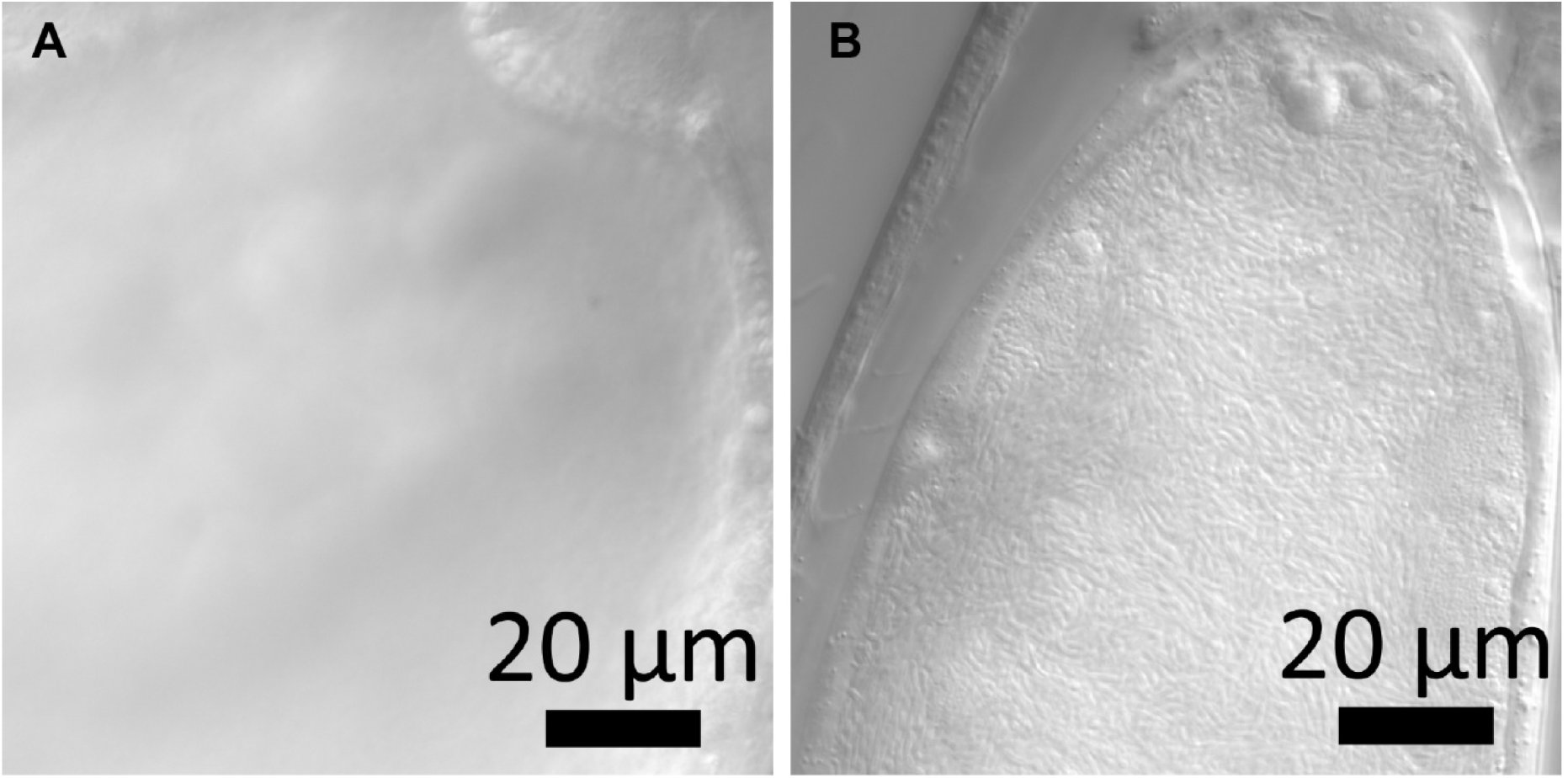
Examining *S. hermaphroditum* intestines for the presence of bacteria. A) The anterior intestine of an adult *S. hermaphroditum* hermaphrodite, five days after it was administered to a *Galleria mellonella* larva as an axenically raised IJ. No bacteria can be seen in the intestine in any focal plane, and the lumen of the intestine cannot be easily seen. B) The anterior intestine of an adult *S. hermaphroditum* hermaphrodite, five days after it was administered to a *Galleria mellonella* larva carrying its *Xenorhabdus* bacterial symbiont. The lumen of the intestine is distended by a massive amount of bacteria. Scale bars, 20 µm.

**Figure S3.**
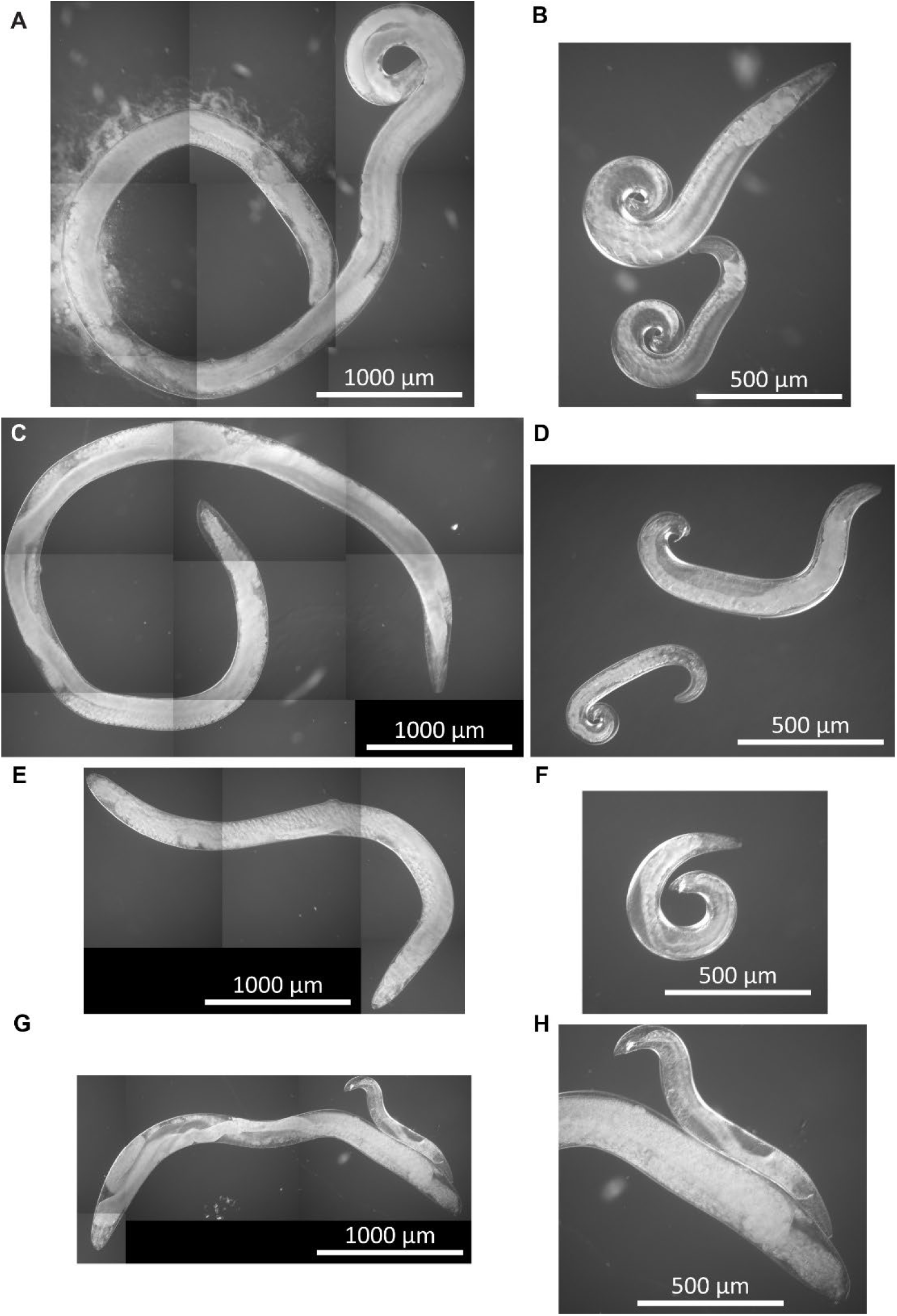
Growth of other *Steinernema* species in living waxmoth larvae. Adult *Steinernema* females and males were taken from living *Galleria mellonella* larvae, five days after the administration of asymbiotic IJs. A) A *Steinernema carpocapsae* (Italy) female. B) Two *Steinernema carpocapsae* (Italy) males. C) A *Steinernema carpocapsae* (Poland) female. D) Two *Steinernema carpocapsae* (Poland) males. E) A *Steinernema feltiae* female. F) A *Steinernema feltiae* male. G) A *Steinernema glaseri* female and male. H) Magnified view of the *Steinernema glaseri* male in panel G. Scale bars as indicated; panels A, C, E, and F are at the same magnification, and panels B, D, F, and H are each at twice the magnification of panel A.

**Figure S4.**
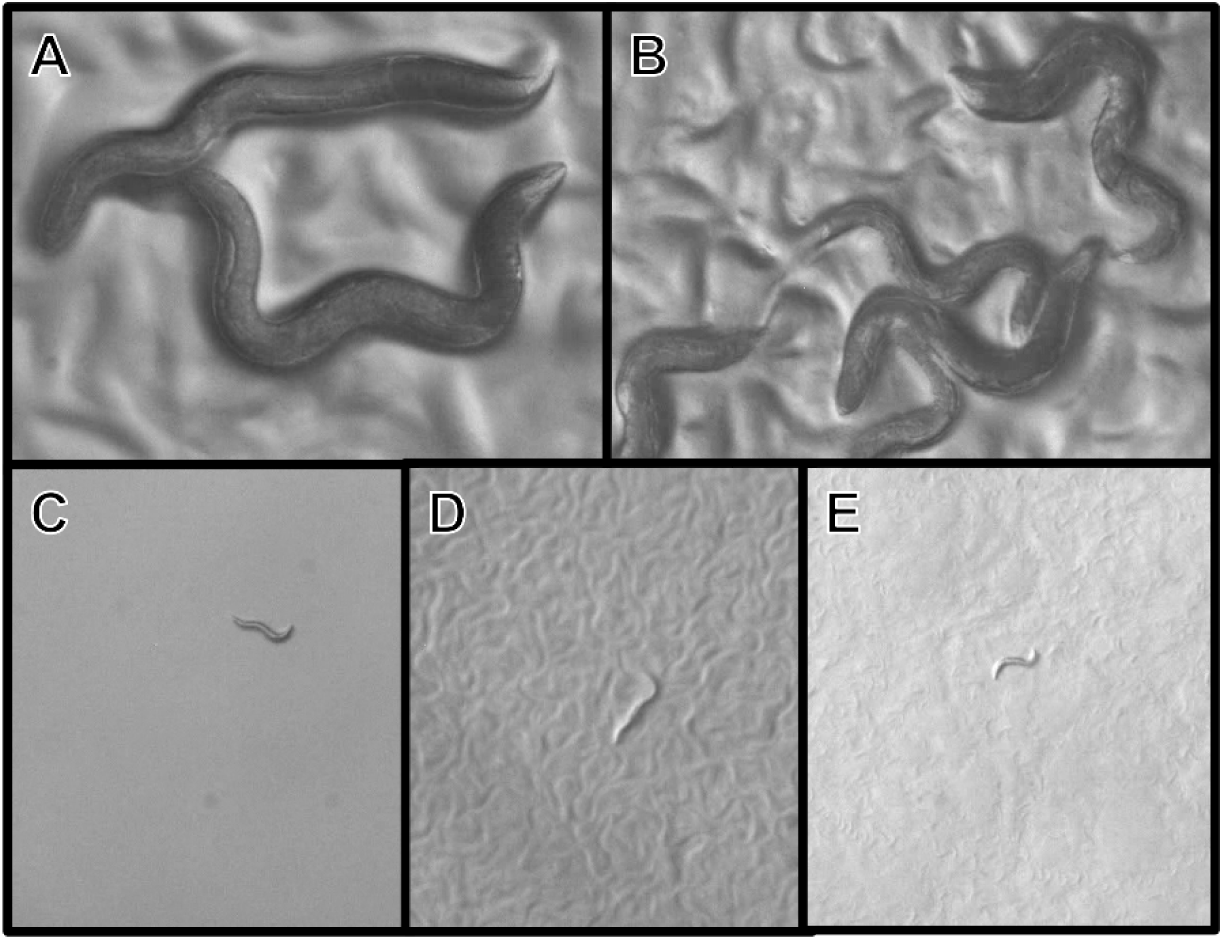
Growth of *S. hermaphroditum* on the waxworm-derived strain of *Serratia marcescens*. Images of *S. hermaphroditum* animals three days after their eggs were placed on different combinations of eggs and media and incubated at 25°C. All images are stills from videos taken at the same magnification. A) Incubated on *Comamonas aquatica* DA1877 on EPM. B) Incubated on *Xenorhabdus griffiniae* on NGM. C) Incubated on NGM without bacteria. D) Incubated on the *Serratia marcescens* isolate on EPM. E) Incubated on the *Serratia marcescens* isolate on EPM.

**Figure S5.**
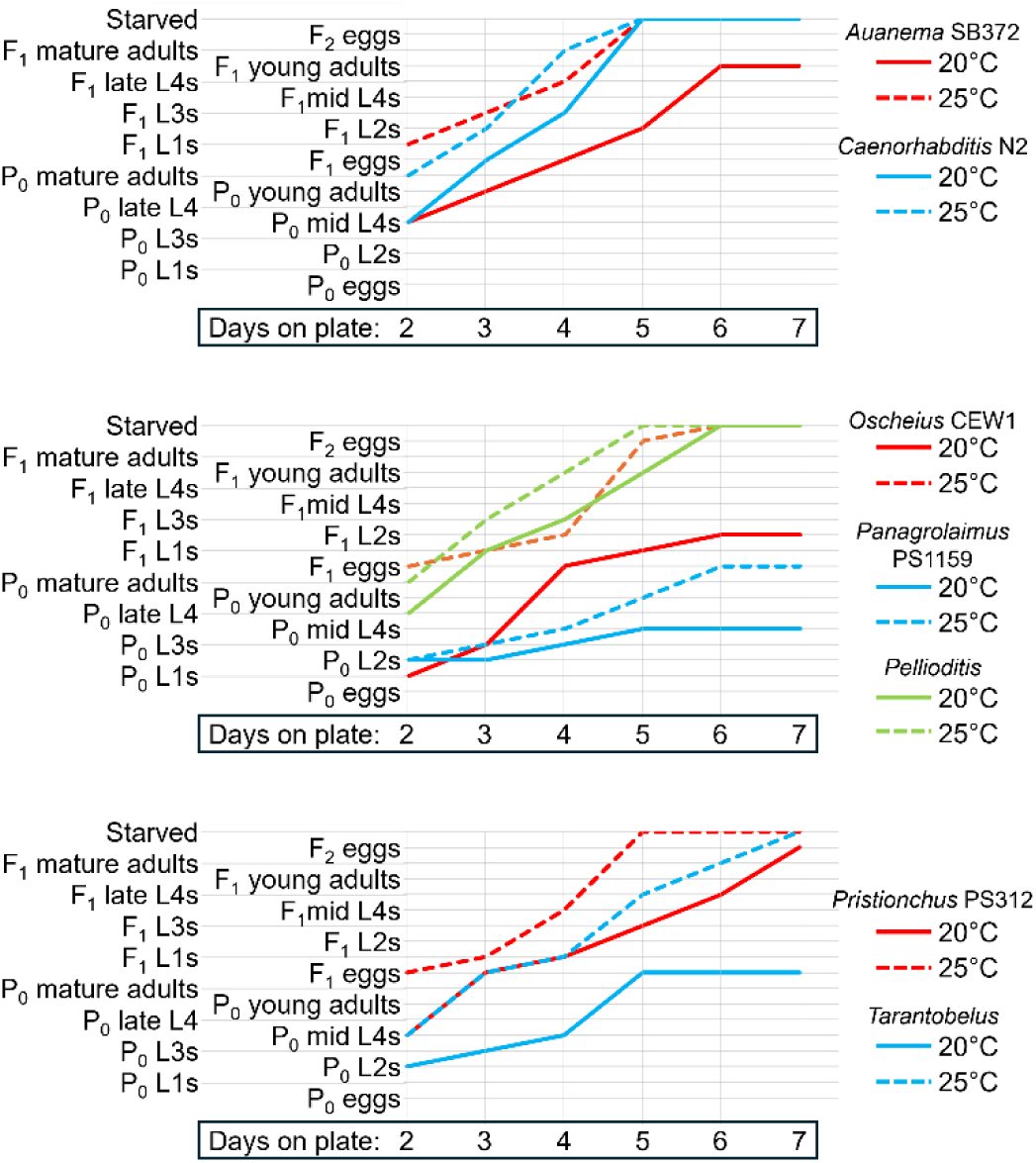
Growth curves of populations of different species at 20°C and at 25°C. Axenized eggs were placed on lawns of *E. coli* OP50 bacteria on NGM agar media and grown at the indicated temperature for seven days. The plates were examined each day, and populations were staged based on morphological markers.

**Video S1 A population of *S. hermaphroditum* discovered in a living *Galleria mellonella* waxmoth larva**. This video was taken six days after six IJs were pipetted onto the back of the insect. The insect can seen to be moving at the start of the video, and after the insect is bisected a mixed-stage population of worms can be seen to spill forth.

**Video S2 Dissection of a living waxmoth larva shows adults growing in the hemolymph under the cuticle.** Four days after the administration of IJs, a living waxmoth larva was first decapitated and then its cuticle was cut open along the ventral midline, with the animal lying on its back. Large adult worms can be seen swimming in the hemolymph just under the cuticle, with intact tissues beneath them.

